# Adenylyl cyclases combinatorially integrate opposing dopamine receptor signals

**DOI:** 10.64898/2026.07.10.737756

**Authors:** Jan Gregrowicz, Michael B. Elowitz

**Affiliations:** Division of Biology and Biological Engineering, California Institute of Technology, Pasadena, CA 91125, USA; Howard Hughes Medical Institute, California Institute of Technology, Pasadena, CA 91125, USA

## Abstract

Dopamine receptors are divided into two families which exert opposing effects on the second messenger cyclic AMP (cAMP). While most neuronal cell types express a single receptor subtype, some neurons co-express opposing receptor subtypes. It remains unclear how these cells could resolve simultaneous stimulatory and inhibitory inputs. Here, we introduce a multiplexed assay that quantifies surface receptor abundance and dynamic cAMP output in single cells. Using this assay, together with mathematical modeling, we demonstrate that signals from opposing receptor subtypes are integrated flexibly by downstream adenylyl cyclases (ACs) rather than at the receptor level. Because AC isoforms exhibit unique biochemical properties, a cell’s AC expression profile determines whether conflicting inputs are cancelled, suppressed, or amplified. Brain transcriptome analysis indicates that co-expression of opposing dopamine receptors is associated with expression of specific AC isoforms predicted to sustain signaling during multi-receptor activation. Our results show that dopamine signal integration depends on the expression profiles of receptors and AC isoforms in a predictable way.

## Introduction

Dopamine is a neuromodulator that governs learning, motivation, motor control, and hormone release. Its dysregulation underlies disorders ranging from Parkinson’s disease to schizophrenia and addiction^1–3^. Its diverse physiological effects are mediated by five G protein-coupled receptor (GPCR) subtypes (DRD1 through DRD5). These receptors are classically grouped into two families with opposing downstream effects: D1-like receptors (DRD1, DRD5) couple to Gαs/olf proteins to stimulate cAMP production, whereas D2-like receptors (DRD2, DRD3, DRD4) couple to Gi/o proteins to inhibit cAMP production^4^.

Canonical models of dopamine signaling assume that distinct neuronal populations express either activating or inhibitory receptors to drive opposing programs. A classic example is the segregation of D1-expressing and D2-expressing medium spiny neurons (MSNs) of the direct and indirect pathways in the basal ganglia^5,6^ (**Fig. 1)**. However, evidence from single-cell RT-PCR^7^, single-cell transcriptomics^8^, and transgenic reporters^9^ indicates that subpopulations of neurons frequently co-express receptor subtypes from opposing families. Although rare, representing between 6-16% of the basal ganglia neurons^10^, these co-expressing populations form distinct projections and modulate functionally relevant outputs^9,10^. These co-expressing populations appear to serve a computational function: in Drosophila reward neurons, individual presynaptic terminals co-express stimulatory DRD1 and inhibitory DRD2 to bidirectionally encode specific reward intensities^11^. The presence of this dual-receptor architecture across vertebrate and invertebrate systems suggests it is an evolutionarily conserved signal-processing strategy.

**Figure 1.**
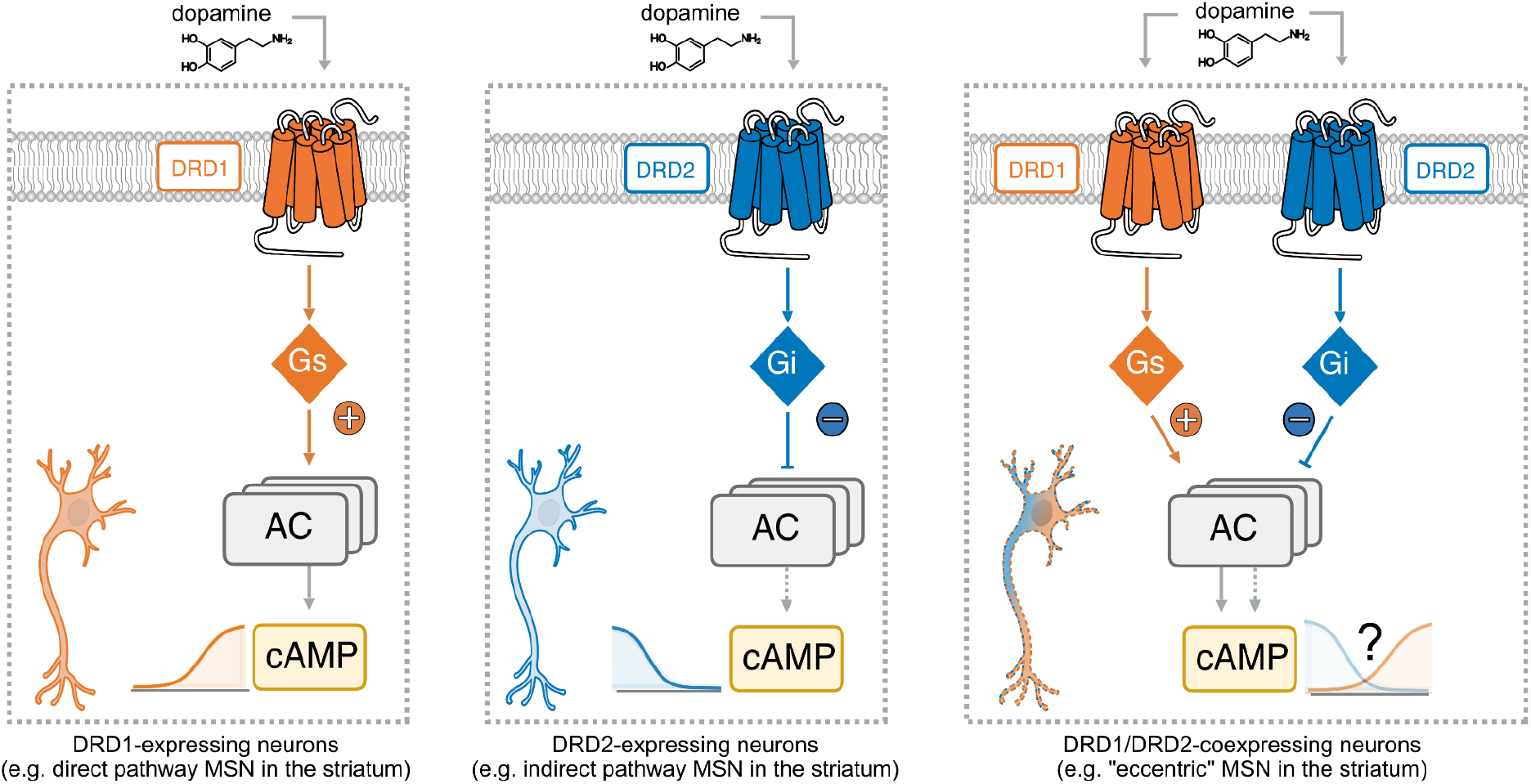
Dopamine signaling in cells expressing DRD1, DRD2, or both. Schematic of three classes of dopamine-responsive neurons. (Left) DRD1-expressing neurons, exemplified by direct-pathway medium spiny neurons (dMSNs) in the striatum, couple dopamine to Gαs-mediated activation of a set of adenylyl cyclase variants, which drive cAMP production. (Middle) DRD2-expressing neurons, exemplified by indirect-pathway MSNs (iMSNs) in the striatum, couple dopamine to Gαi-mediated inhibition of AC, suppressing cAMP production. (Right) A minority (6-16%) of “eccentric” MSNs (eMSNs) in the striatum co-express both DRD1 and DRD2, simultaneously engaging Gαs and Gαi on a shared AC pool. The resulting cAMP response, and the mechanisms that shape it, remain poorly understood.

Several mechanisms could in principle generate the distinct dopamine responses observed in cells expressing D1, D2, or both receptors. Differences in receptor affinity have been proposed to temporally segregate tonic (low) and phasic (high) dopamine signals, with the high-affinity DRD2 receptor detecting baseline tonic dopamine and the lower-affinity DRD1 engaging only during phasic bursts^12^. However, this temporal segregation does not resolve the integration problem in co-expressing cells: during phasic bursts, DRD1 engages while DRD2 remains near-saturated by tonic baseline, so both pathways must be integrated simultaneously. Receptor heteromerization provides an alternative explanation: D1-D2 complexes have been reported to switch coupling from canonical Gαs and Gαi to Gq/11^13,14^, although the physiological relevance of these complexes remains debated^15^. Beyond these receptor-centric mechanisms, expression-level variation in receptors, transducers (Gαs, Gαi, Gβγ), and downstream effectors such as adenylyl cyclases could each independently shape the integrated response. This raises a fundamental question: how do co-expressed dopamine receptors combine to integrate dopamine inputs, and how does signal integration vary among cell types expressing different levels of dopamine pathway components?

Quantitatively dissecting how D1 and D2 co-expression shapes the dopamine response requires explicit measurement of receptor levels, control of transducer availability, and quantification of signaling output. Such measurements, however, have proven difficult in practice. Most experiments rely on cellular backgrounds in which receptor abundance, surface delivery, and transducer stoichiometry are neither quantified nor independently controlled^16^. Because receptor expression levels alone can shift apparent potency, basal activity, and even qualitative agonist behavior^17,18^, it is often impossible to disentangle intrinsic receptor properties from variation in expression.

Here, we combine a quantitative multiplexed single-cell signaling assay with mechanistic modeling to establish a predictive framework for dopamine signal integration. By simultaneously quantifying surface receptor abundance and cAMP output in the same cells **(Fig. 2)**, we show that single-receptor behavior is sensitive to both receptor dosage and G-protein availability **(Fig. 3)**. Building on these single-receptor measurements, we then ask whether pairwise receptor combinations can be predicted from individual receptor behavior. Gαs-coupled receptors systematically dominate over Gαi-coupled receptors to a degree that simple additive models cannot account for **(Fig. 4)**. We trace this discrepancy to the adenylyl cyclase (AC) layer: endogenous AC isoforms exhibit distinct expression profiles, and differ in their sensitivity to Gαi inhibition, their potentiation by Gβγ, and their basal catalytic activity. Incorporating these features in the model explains responses in cells expressing multiple dopamine receptors and diverse AC profiles **(Fig. 5)**. Finally, the model suggests an explanation for an association between AC2 expression and co-expression of opposite-acting dopamine receptors, which we identified in brain atlas data sets (**Fig. 6**). Based on these results, we propose that cell-type-specific AC transcriptomic programs represent a general determinant of how neuromodulatory inputs are transformed into cellular outputs.

**Figure 2.**
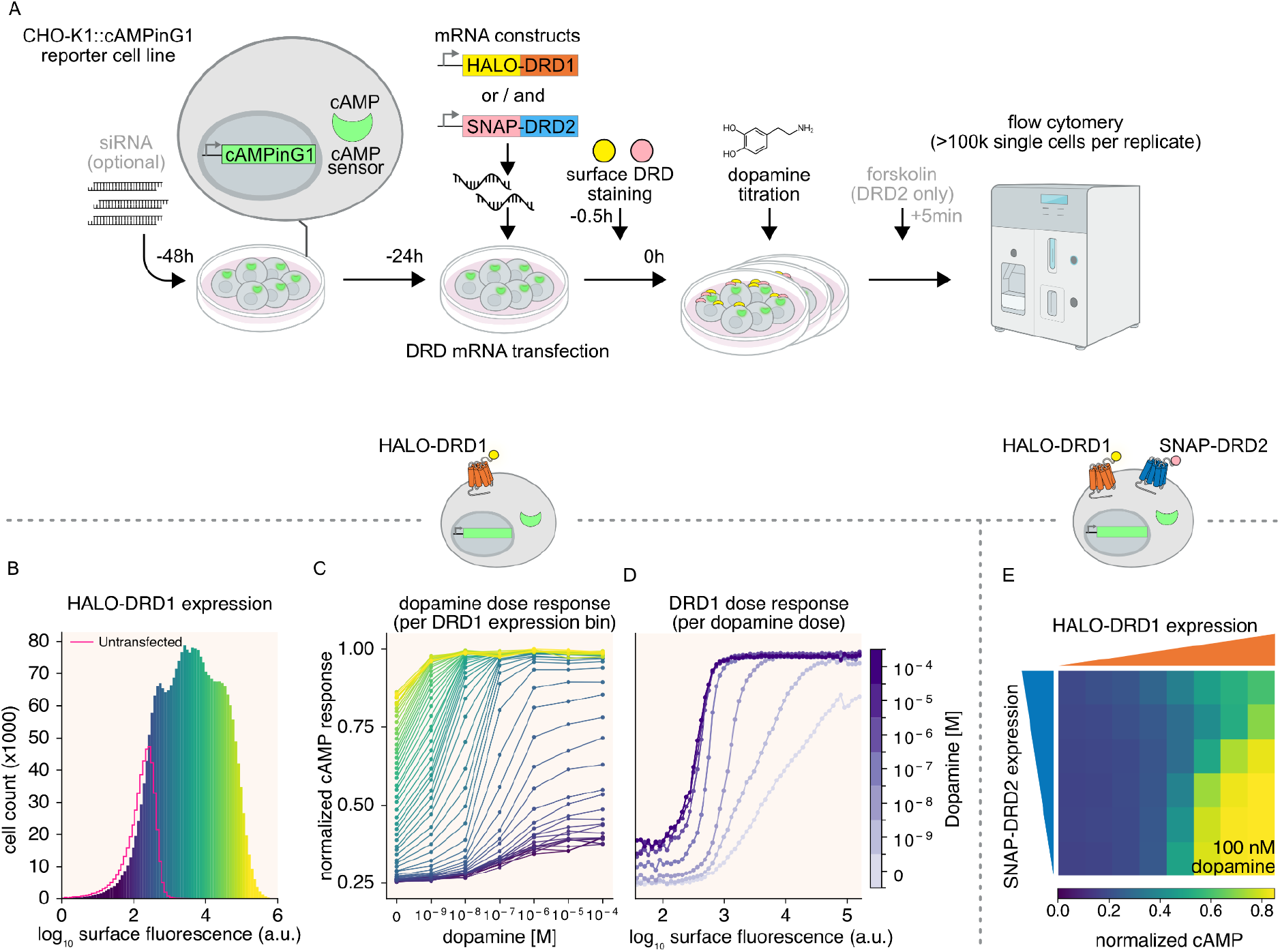
Single-cell quantification of surface dopamine receptor abundance and dopamine-evoked cAMP. (A) Experimental workflow. A stable monoclonal CHO-K1 line expressing the ratiometric cAMP biosensor cAMPinG1 (CHO-K1::cAMPinG1) is transfected with mRNA encoding HALO-DRD1 and/or SNAP-DRD2 24 h before the assay. Optional siRNA transfection (48 h before the assay) enables knockdown of downstream signaling components. Surface receptors are labeled with cell-impermeable HALO- and SNAP-ligands 30 min before dopamine titration. For DRD2-only conditions, forskolin is added 5 min before flow cytometry readout (>100,000 single cells per replicate). (B) Single-cell distribution of surface HALO-DRD1 expression after mRNA transfection (colored histogram); pink line shows untransfected control. mRNA transfection yields a broad, continuous range of receptor abundances. Color bins are reused in panel C. (C) Dopamine dose-response curves binned by DRD1 surface expression (colors as in B). Higher surface DRD1 amplifies both the maximal cAMP response and apparent dopamine sensitivity. (D) cAMP response as a function of surface DRD1 expression at fixed dopamine concentrations (line color encodes dopamine dose; see colorbar). At zero dopamine, the curve reveals constitutive DRD1 signaling. At high dopamine, the response becomes ultrasensitive to surface receptor abundance, so small decreases in surface DRD1 abolish responsiveness. (E) Heatmap of normalized cAMP response to 100 nM dopamine in cells co-expressing HALO-DRD1 (x-axis) and SNAP-DRD2 (y-axis). The cAMP output is jointly determined by the ratios of both receptors at a fixed dopamine dose.

**Figure 3.**
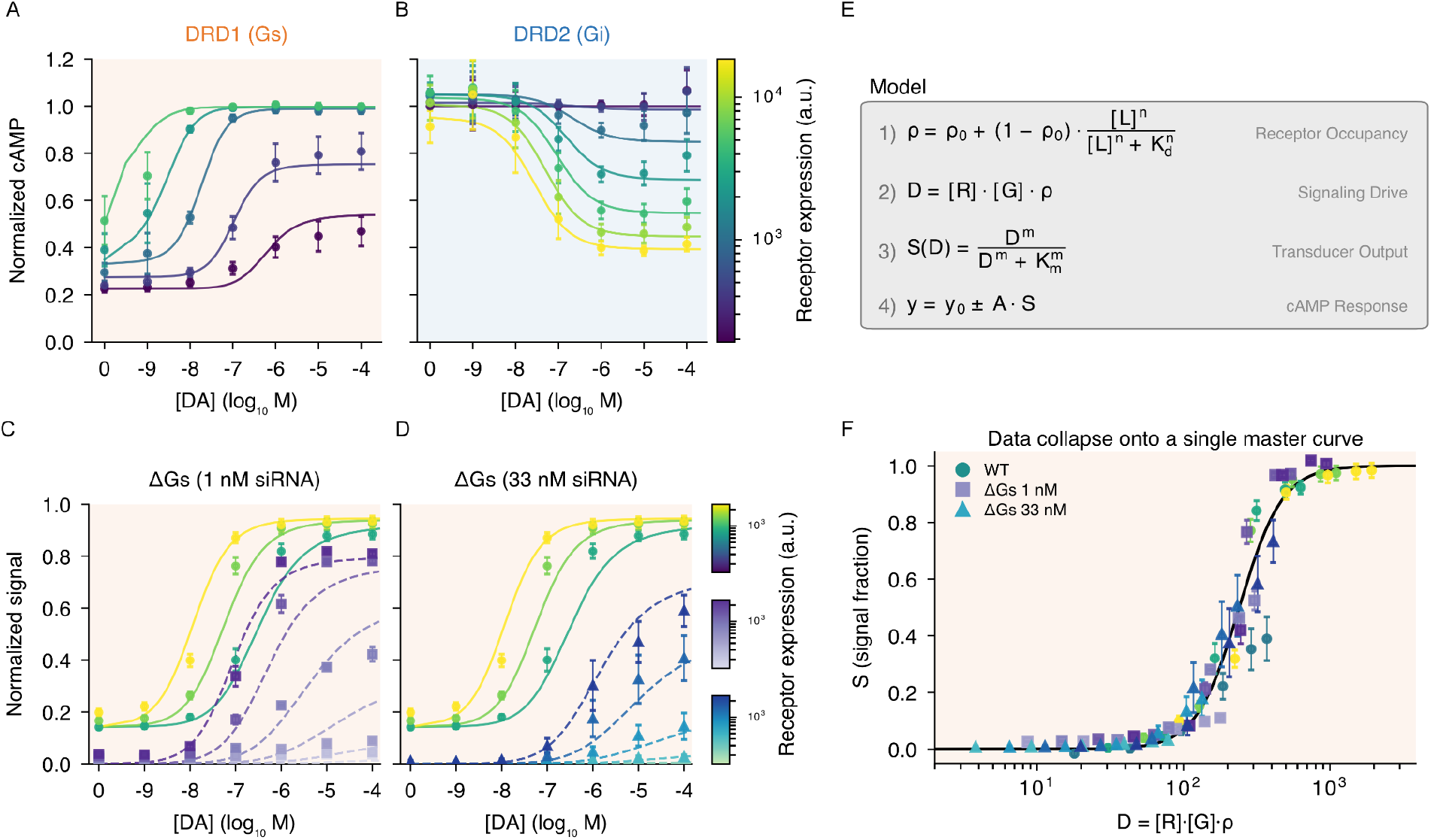
A unified operational model captures receptor-dependent dopamine signal integration. (A, B). Single-cell dose-response curves for (A) Gαs-coupled DRD1 and (B) Gαi-coupled DRD2 in CHO-K1::cAMPinG1 cells transiently expressing HALO-tagged receptors. DRD2 was stimulated in the presence of 10 µM forskolin to elevate the baseline cAMP level. Cells are binned and colored by their measured surface receptor abundance (arbitrary flow-cytometry units; see color bars). Solid lines represent joint fits of the operational model (R^2^ = 0.98 for DRD1; R^2^ = 0.97 for DRD2) with binding affinity (Kd) locked to established IUPHAR values (fit values and 95% CIs are provided in Table S1). Data represent per-bin medians, with error bars indicating ±1 SEM across independent replicate dates (n = 4 for DRD1, n = 5 for DRD2). The remaining subtypes (DRD3, DRD4, DRD5) are characterized in Fig. S3. (C, D) DRD1 dopamine dose-response under siRNA knockdown of Gαs at (C) 1 nM and (D) 33 nM siGNAS (ΔGs), overlaid on the wild-type Gαs response, binned by surface DRD1 abundance (colors as in A). Solid lines are the shared-parameter operational-model fit in which only the available transducer pool [Gs] varies per condition (Table S2); Gαs depletion right-shifts the curves along the dose axis without lowering the maximal response. (E). Schematic of the modified operational model. Ligand binding ([L]) yields a fractional receptor occupancy (ρ). This fraction is multiplied by the measured surface receptor abundance ([R]) to yield a composite signaling drive (D = [R] · [G] · ρ). The drive is processed through a saturable Hill-type transducer function (S) to compute the final integrated cAMP output (y). Full mathematical derivation is provided in the Supplementary Information. (F). Master-curve collapse across receptor abundances. Plotting the normalized transducer output (S) against the composite signaling drive (D) collapses the variable dose-response families onto a single, invariant master curve for each receptor subtype. This confirms that cellular cAMP responses are governed by the combined drive quantity rather than receptor density or ligand concentration independently.

**Figure 4.**
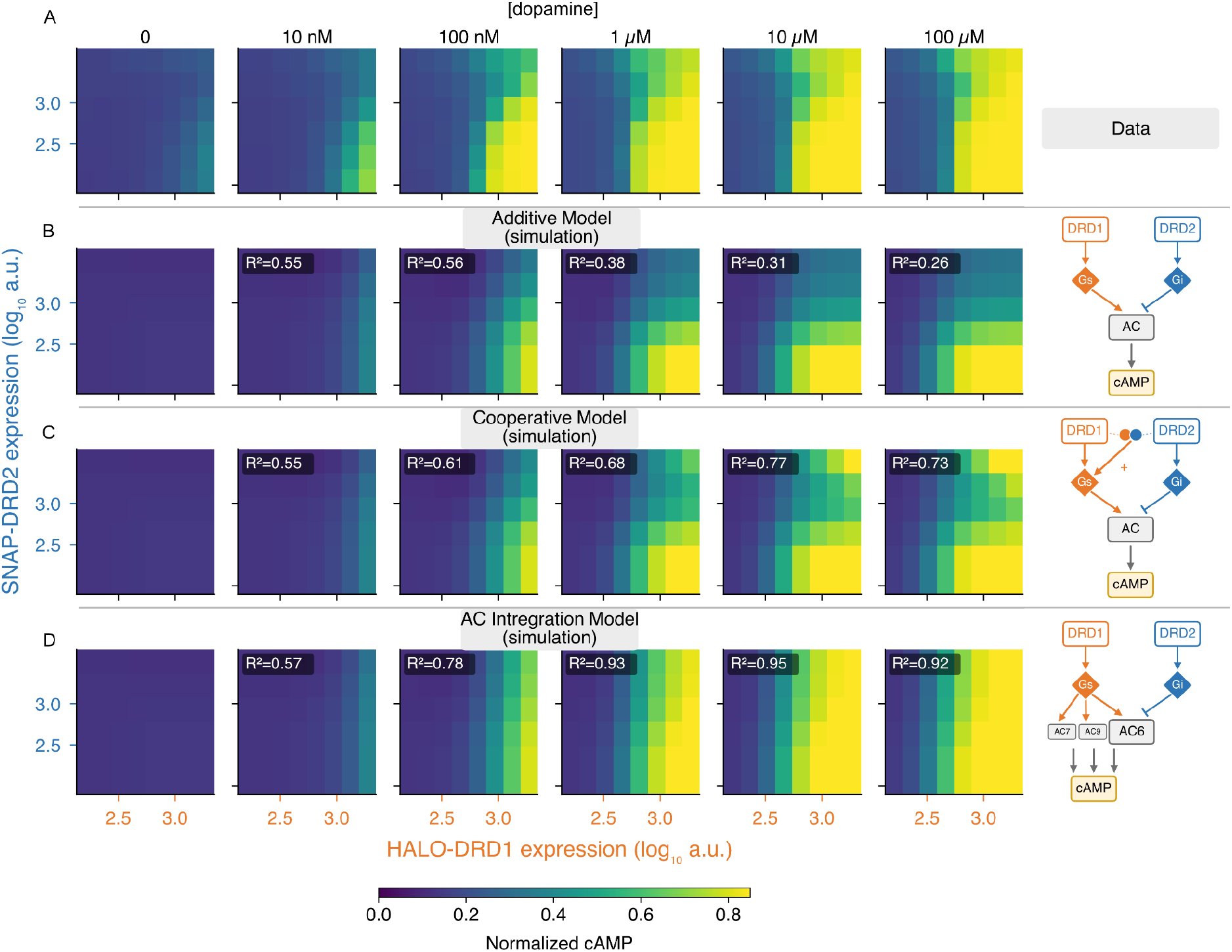
(previous page). Model comparison suggests DRD1–DRD2 signal integration by the adenylyl cyclase layer. CHO-K1 cells co-expressing DRD1 and DRD2 were surface-labeled and binned on a two-dimensional log_10_(DRD1) × log_10_(DRD2) abundance grid. Dopamine dose-responses were reconstructed from single-cell cAMP biosensor readouts (n = 3 biologically independent replicates). **(A)** Two-dimensional cAMP response surfaces across increasing dopamine concentrations. Top row: measured per-bin median normalized cAMP showing competitive cancellation at low concentrations and DRD1 dominance at saturating dopamine concentrations. Bottom row: baseline predictions derived entirely from single-receptor fits assuming independent Gαs and Gαi convergence on a shared adenylyl cyclase pool. While this model reproduces the competitive shape at lower concentrations (10 nM to 100 nM), it systematically under-predicts cAMP in the high-DRD1/high-DRD2 corner at higher dopamine concentrations. **(B)** Competing models for the integration step at saturating dopamine (100 μM). The Additive Model (R^2^ = 0.55) assumes a homogeneous cyclase pool, predicting a mutual cancellation that fails to capture the measured response. **(C)** The Cooperative Model (R^2^ = 0.75) incorporates a single-parameter cooperative term representing receptor-level crosstalk due to heterodimerization to partially recapitulate signaling profiles at high dopamine. **(D)** The AC Integration Model (R^2^ = 0.87 globally) explicitly models three endogenously expressed adenylyl cyclase isoforms (AC6, AC7, AC9) in their measured reporter-line proportions. Signal-flow diagrams (bottom) illustrate this distinction: modeling AC6 as an integrator of Gαs and Gαi, and AC7 and AC9 as strictly Gαs-responsive. Model selection favored the AC Integration Model by AICc (ΔAICc = 465 versus the additive model, 247 versus the cooperative model) and by held-out R^2^ (0.63, 0.77, and 0.88 for the additive, cooperative, and AC Integration models, respectively).

**Figure 5.**
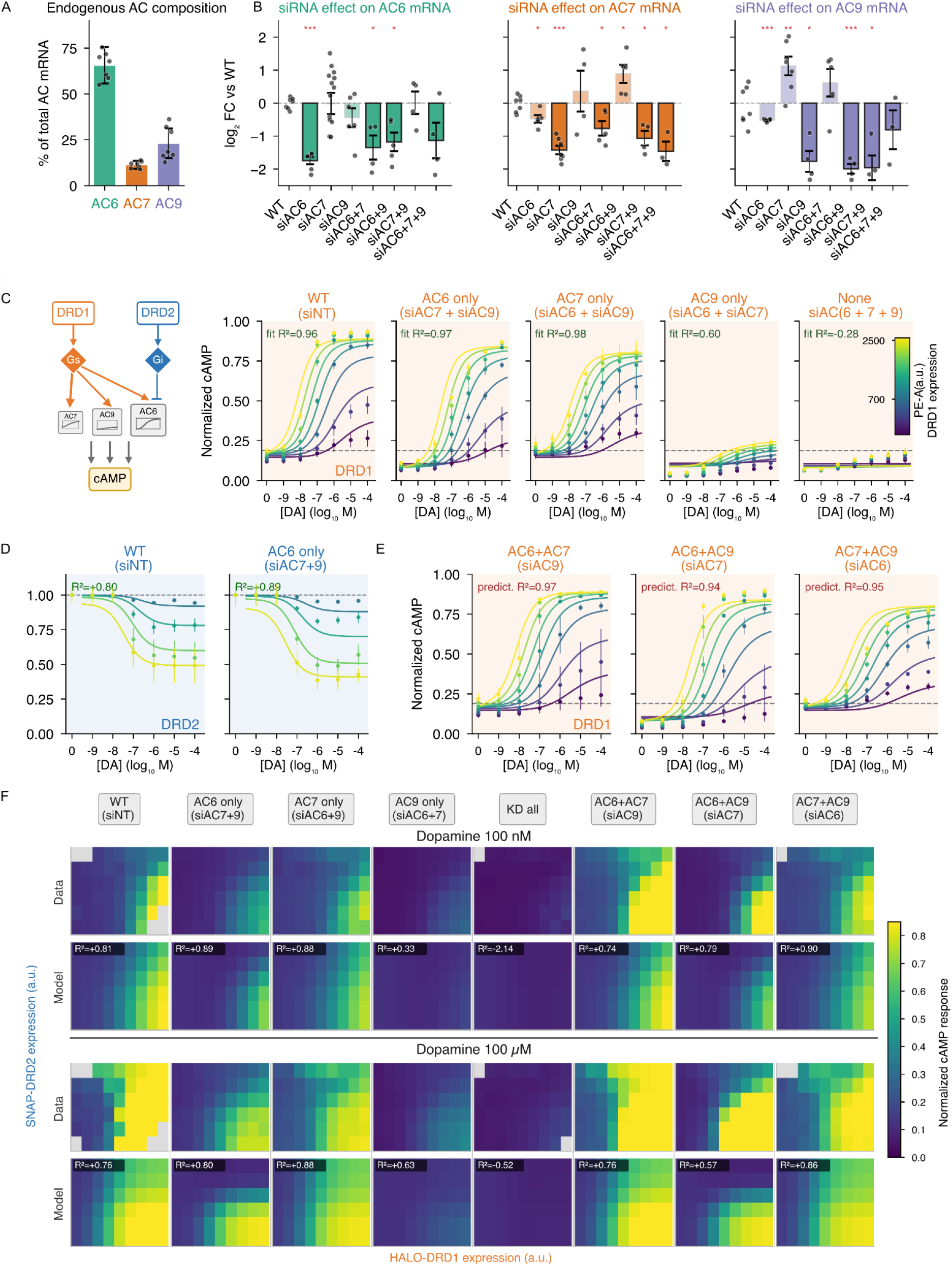
(previous page) Adenylyl cyclase composition predicts dual-receptor signal integration. **(A) Endogenous adenylyl cyclase composition**. Baseline mRNA fractions of AC6, AC7, and AC9 in the CHO-K1 reporter line (AC6 65.6 ± 10.0%, AC7 11.3 ± 2.2%, AC9 23.1 ± 8.1%; n = 7 runs; dots, individual runs). **(B) siRNA knockdown efficiency**. Log_2_ fold change of AC transcripts following single, double, or triple siRNA depletion. Bold borders indicate targeted genes. Data are mean ± SEM (dots represent independent qPCR runs; n = 3–14). Significance was determined by a one-sample, two-tailed t-test on log_2_ fold change versus 0: *p < 0.05, **p < 0.01, ***p < 0.001. **(C) Extracting activation parameters in isolated AC contexts**. Left schematic illustrates the AC-layer wiring for DRD1 (Gαs) and DRD2 (Gαi/Gβγ). Right subpanels show single-cell DRD1-mediated cAMP dose-response curves across varying surface receptor abundances (color scale, 700 to 2500 a.u.). Solid lines indicate operational-model fits derived exclusively from these isolated single-AC contexts. Error bars represent ±1 SEM across independent experimental dates (n = 3–4 replicates per condition). **(D) AC6 dictates the DRD2-mediated Gαi inhibitory response**. Dose-response curves of DRD2 stimulated against a 10 µM forskolin baseline in wild-type and AC6-only contexts. Solid lines represent the joint model fits for Gαi-mediated inhibition. Error bars represent ±1 SEM (n = 3–4 replicates). **(E) Parameter-free predictions of dual-isoform mixtures**. The single-AC parameters derived in (C) were used to predict DRD1 dose-response curves in dual-AC contexts. Solid curves represent transcript-weighted linear combinations of the isolated parameters, generated without additional fitting. Error bars represent ±1 SEM (n = 3 replicates). **(F) Parameter-free prediction of the full combinatorial DRD1 × DRD2 response across adenylyl cyclase contexts**. Single cells co-expressing HALO-DRD1 and SNAP-DRD2 were binned on a two-dimensional surface-abundance grid (9 HALO-DRD1 × 6 SNAP-DRD2 bins) and stimulated with 100 nM or 100 µM dopamine across eight AC-knockdown contexts: WT (siNT), AC6 only (siAC7+9), AC7 only (siAC6+9), AC9 only (siAC6+7), AC6+AC7 (siAC9), AC6+AC9 (siAC7), AC7+AC9 (siAC6), and triple knockdown (siAC6+7+9). For each dopamine concentration, the upper row shows the measured per-bin median normalized cAMP (Data) and the lower row shows the prediction of the Isoform-resolved AC Integration Model (Model), evaluated with all parameters fixed from the preceding panels and no refitting to these data. Color encodes normalized cAMP (contamination-corrected and reporter-background-subtracted). Per-bin values are medians of single cells pooled across independent experiments; grey bins denote fewer than 50 cells and were excluded. Per-panel R^2^ (model versus data) is annotated on each tile; the model reproduces the measured landscapes across all contexts (pooled global R^2^ = 0.876 across 847 bins).

**Figure 6.**
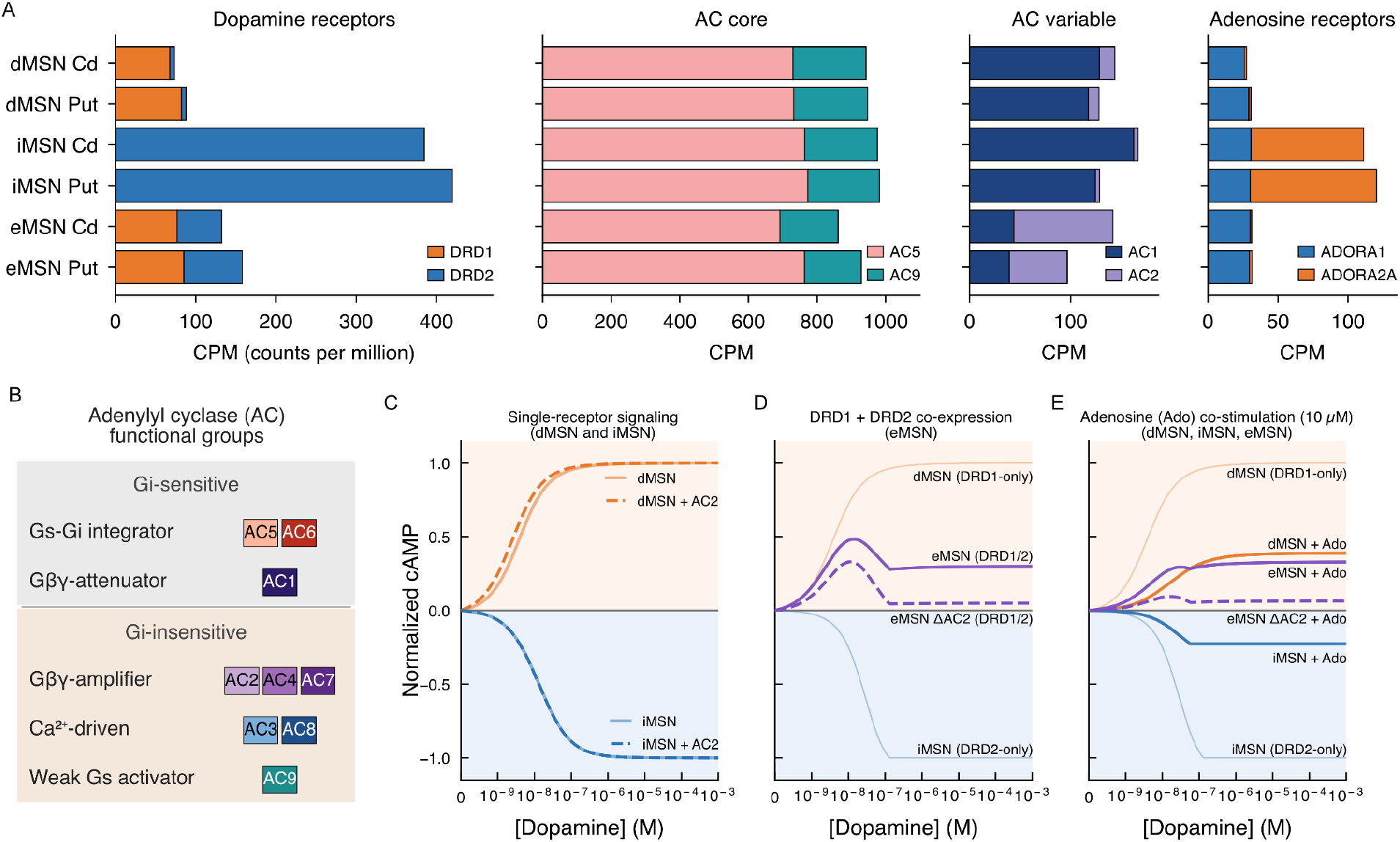
AC composition predicts cell-type-specific dopamine integration in striatal neurons. **(A)** Cell-type specific transcriptomic quantification of dopamine receptors, the adenylyl cyclase (AC) core, the variable AC pool, and adenosine receptors across striatal dMSNs, iMSNs, and eMSNs. While dMSNs and iMSNs rely on a core of AC5 and AC9 alongside the variable isoform AC1, eMSNs specifically upregulate AC2. **(B)** A biochemical dictionary of adenylyl cyclase integration. Isoforms are defined by their established regulatory logic: AC5 integrates opposing Gαs and Gαi signals, AC9 is fundamentally Gαi-insensitive, and AC2 acts as a conditional amplifier potentiated by Gβγ during concurrent Gαi activation. **(C)** Operational model simulations of single-receptor-dominated MSNs (dopamine only). The hypothetical addition of AC2 (dotted line) does not alter the dopamine response in strictly DRD1- or DRD2-dominated cells. **(E)** eMSN simulated dopamine integration (dopamine only). In a hypothetical profile matching canonical MSNs (AC2 reduced to dMSN level, dotted line), concurrent DRD1 and DRD2 activation results in signal collapse. In the WT eMSN profile (solid line), the conditional Gβγ amplifier AC2 buffers the amplitude, capturing an initial DRD1-driven peak before sustaining a lower, graded plateau. **(E)** Simulated co-stimulation of dopamine and adenosine. Under concurrent stimulation, iMSNs exhibit a high baseline that is canceled by increasing dopamine. In eMSNs, a hypothetical, reduced AC2 background results in a signal collapse toward baseline whereas the WT eMSN profile synergistically scales, allowing the co-expressing eMSN to converge with the robust dMSN signaling profile.

## Results

### Simultaneous quantification of surface receptors and cAMP captures single-cell signaling profiles

The dopamine signaling pathway maps dopamine inputs onto cAMP outputs. Understanding this quantitative input-output relationship is difficult with bulk measurements because they average over cell-cell variation in the expression levels of receptors and other components. We therefore developed a single-cell assay that simultaneously measures surface receptor density and cAMP responses to varying dopamine concentrations (**Fig. 2A**).

To quantify cAMP levels, we engineered a stable monoclonal cell line (CHO-K1::cAMPinG1) expressing the ratiometric cAMP sensor cAMPinG1 (**Fig. 2A, Methods**). Systematically mapping combinatorial signaling requires testing many perturbation conditions across hundreds of thousands of individual cells, a scale that necessitates tractable and transfectable *in vitro* models. We used CHO-K1 cells as a base cell line because they lack endogenous dopamine receptors (reducing background signaling) but otherwise express all necessary cAMP transduction machinery (Gαs, Gαi1,2,3, and adenylyl cyclases AC6, AC7, AC9) (**Fig. S1**). These endogenous CHO transducers and cyclases share conserved biochemical regulatory logic with the isoforms expressed in mammalian neurons, providing a functionally homologous background for studying signal integration. Importantly, the assay also allows for optional siRNA transfection to selectively knock down specific downstream transducers and adenylyl cyclases alone or in combinations.

To measure dopamine receptor expression in individual cells, we introduced N-terminally tagged (SNAP or HALO) dopamine receptors via mRNA transfection. The cell-impermeable fluorescent HALO- and SNAP-ligands ensure that only the actively signaling, surface-localized receptor pool was quantified. mRNA transfections yielded uniform expression distributions, allowing us to sample a broad, continuous range of receptor abundances (**Fig. 2B**).

Using this assay, we first constructed a family of dopamine-cAMP dose-response curves for varying expression levels of the Gs-coupled DRD1 receptor. As expected, higher surface receptor density amplified both the absolute cAMP response and the sensitivity to dopamine (**Fig. 2C**). Conversely, plotting cAMP output against receptor abundance at fixed ligand concentration allowed measurement of basal signaling by DRD1 in the absence of dopamine (**Fig. 2D)**. It also showed that, at higher dopamine levels, signaling becomes ultrasensitive to DRD1 expression levels. In this regime, removing even a small fraction of receptors (e.g., through desensitization) is sufficient to render an otherwise dopamine-responsive cell unresponsive.

To understand how multiple receptors integrate dopamine signals, we co-expressed functionally opposed DRD1 and DRD2 receptors. The two receptors were labeled, respectively, with orthogonal SNAP and HALO tags to allow simultaneous quantification of surface DRD1 and DRD2 abundance within the same cell along with the cAMP dopamine response (**Fig. 2E**). To illustrate this multi-dimensional space, the heatmap shows a response to 100 nM dopamine, demonstrating how the ratio of the two receptors dictates cAMP levels at a fixed dopamine dose.

We used a high-throughput format to analyze many different receptor and dopamine conditions in parallel over time. To avoid time-dependent drift in measurements, all assays were performed in the presence of the phosphodiesterase inhibitor IBMX, which prevents cAMP degradation and allows for sustained signal integration (**Methods**). In addition, to ensure measurements were at steady-state, we performed flow cytometry at slow, but continuous, flow rates. While the rate of cAMP accumulation depended on surface receptor expression **(Fig. S2)**, responses plateaued within a few minutes. Based on this analysis, we made endpoint measurements at 5 minutes after dopamine addition.

### Receptor and G-protein availability combinatorially dictate single-receptor dopamine responses

Having established the assay, we next validated signaling by DRD1 and DRD2. Dopamine stimulation drove cAMP accumulation through the Gαs-coupled DRD1 (**Fig. 3A**) and suppressed forskolin-elevated cAMP through the Gαi-coupled DRD2 (**Fig. 3B**). We also characterized the remaining subtypes (DRD3, DRD4, DRD5) **(Fig. S3**).

For both DRD1 and DRD2, increasing receptor abundance shifted the dopamine-cAMP input-output curves toward lower dopamine concentrations (increasing sensitivity) in a receptor level-dependent way (**Fig. 3A,B**). It also shifted the amplitude of response upward for DRD1 (greater stimulation) and downward for DRD2 (greater dopamine-induced inhibition of the forskolin-elevated baseline).

This dual scaling can be explained by a modified operational model (**Fig. 3E**; derivation in SI). In this model, the cAMP response depends on a composite signaling drive D = [R]·[G]·ρ([L]), which combines surface receptor abundance [R], available transducer pool [G], and the dopamine-bound fraction of receptor ρ. The drive feeds into a Hill-type transducer function S(D) with half-saturation Km and cooperativity m, and the cAMP response is y = y_0_ ± A·S (addition for Gαs; subtraction for Gαi). At low receptor abundance, D remains sub-saturating (D < Km) even when receptor occupancy is complete, so S falls short of 1 and the response does not reach its maximum amplitude, A. Fixing the dopamine binding affinities (Kd) to IUPHAR database values, a single set of receptor-specific parameters was able to fit the data for each subtype (numerical values in SI; R^2^ = 0.98 for DRD1, 0.97 for DRD2).

We next asked whether the model could capture the system’s dependence on G-protein levels. To test this, we used siRNA to deplete Gαs. At both 1 nM and 33 nM siRNA doses, Gαs depletion systematically right-shifted the DRD1 dose-response curves toward higher dopamine concentrations across all receptor abundance levels (**Figure 3C, D**). Our model predicts that reducing the available G-protein pool would produce this shift (Methods; Appendix 1). By fitting the model to these data, we estimated the functional reduction in the Gαs protein pool to be 50% (at 1 nM siRNA) and 79% (at 33 nM siRNA). While qPCR analysis indicated a 93% depletion of Gαs mRNA transcripts (**Fig. S4**), the model provides an estimate of the functional protein pool remaining at the time of the assay. This discrepancy between transcript and functional protein levels could arise from several factors, such as the slower turnover rate of pre-existing Gαs proteins.

Consistent with this prediction, plotting the normalized signal fraction S against the composite signaling drive D = [R]·[G]·ρ([L]) collapses the wildtype, ΔGs 1 nM, and ΔGs 33 nM DRD1 data onto a single curve spanning several orders of magnitude in D (**Fig. 3F**). Physically, [R]·ρ represents the amount of dopamine-bound receptor and [G] is the available transducer pool. The collapse shows that the cAMP response depends only on this combined quantity, not on each factor independently, and that the receptor’s intrinsic parameters (n, m, Km, A) are conserved across cellular contexts.

### Co-expressed DRD1 and DRD2 signal non-additively

Having established a quantitative model of single-receptor responses (**Fig. 3**), we next asked how co-expressed receptors combinatorially shape signaling. We co-transfected CHO-K1 cells with tagged DRD1 and DRD2, and measured integrated cAMP output across a two-dimensional grid of receptor co-expression levels, as in **Fig. 2E**.

At low dopamine (10–100 nM), increasing DRD2 expression reduced the apparent activity of DRD1, consistent with the opposing effects of the two receptors on adenylyl cyclases (ACs) (**Fig. 4A**). At 100 nM, the response was set by the balance of the two receptors, with a diagonal boundary across the DRD1–DRD2 expression plane separating DRD1-driven cAMP from DRD2-suppressed output. However, some DRD1-dependent signaling remained even at the highest DRD2 levels. As dopamine increased to 1–100 µM, this pattern faded and DRD1 stimulation dominated across most receptor combinations. These results provide a systematic quantitative analysis of combinatorial DRD1-DRD2 signaling.

A minimal model, in which DRD1 and DRD2, activate and inhibit, respectively, the AC pool was inadequate to explain these data. The additive model assumes that Gαs and Gαi converge independently on a shared AC pool. It predicts that at high expression of both receptors, DRD2-mediated inhibition should cancel or suppress DRD1-mediated stimulation (**Fig. 4A**). The experimental data qualitatively recapitulated this prediction at low dopamine concentrations (10 nM to 100 nM), but deviated at higher, saturating dopamine concentrations (100 μM), where DRD1 dominated (**Fig. 4A**).

One potential, albeit controversial^15,19,20^, feature often invoked to explain non-additive effects in GPCR signaling is cooperativity due to receptor heterodimerization^21^. We therefore extended the model by allowing formation of DRD1-DRD2 heterodimers (Methods; Appendix 1, Model 2). This model partially recapitulated signaling profiles at high dopamine but generated a non-monotonic dependence on DRD2 that diverges from the data (**Fig. 4C**). Because we allowed heterodimers only to potentiate Gαs signaling with a strength that scaled with co-expression of both receptors, while DRD2-mediated inhibition saturates at a fixed ceiling, the synergy term eventually outgrows the inhibition and produces a rebound at the highest co-expression levels.

An alternative hypothesis centers on the co-expression of multiple adenylyl cyclase variants. The parental CHO-K1 endogenous AC pool is dominated by three isoforms: AC6, AC7, and AC9 (**Fig. S1**). In the reporter line, the fractions of these isoforms measured by qPCR are 66% (AC6), 11% (AC7), and 23% (AC9) (Fig. 5A), and these are the values used in the model. AC6 is a classical integrator, stimulated by Gαs, inhibited by Gi, while AC9 is strictly and weakly Gαs-responsive, and AC7 is synergistically potentiated by the Gβγ subunits liberated during Gαi activation^22^. We incorporated these interactions within a broader “AC Integration Model” that coupled DRD1 (via Gαs) and DRD2 (via Gi) to each AC isoform based on known regulatory interactions, transcriptional expression profiles, and competition between Gαs and Gαi for binding AC6 (Methods; Appendix 1, Model 3). The AC Integration Model accurately captured the major features of the combinatorial cAMP response at both low and saturating dopamine (Fig. 4D; R^2^ = 0.87 globally).

### Targeted knockdowns reveal AC variant specificities

The use of multiple AC variants could enable the cell to modulate its response to upstream dopamine signaling. To quantitatively understand signal integration at the AC level, we sought to systematically isolate and study individual ACs and AC combinations using siRNA knockdowns.

First, to establish the baseline AC distribution, we measured AC expression levels in the CHO-K1::cAMPinG1 reporter cell line using quantitative PCR (**Fig. 5A**). As expected, AC6 expression dominated (66% of the total pool), followed by AC9 (23%) and AC7 (11%) (**Fig. 5A**). Within this background, siRNA knockdowns strongly reduced AC expression across all single, pairwise, and triple AC depletions, without altering Gαs expression (**Fig. 5B**). Together, these results established a system for specific, quantitative perturbation and readout of AC expression levels.

To extract the operational parameters for each AC isoform, we measured the dopamine-cAMP dose response in the presence of DRD1 alone, and each individual AC. Ligand-independent basal cAMP signaling depended strongly on AC effector context (**Fig. 5C**). Knockdown of all three AC isoforms eliminated signaling, as expected (**Fig. 5C**, 5th plot). AC9 alone (double knockdown of AC6 and AC7) permitted only weak signaling at high dopamine levels (**Fig. 5C**, 4th plot). By contrast, despite its lower expression level (11% of the AC pool), AC7 alone generated signal amplitudes comparable to the full wild-type expression profile, indicating either high specific activity and/or additional Gβγ-driven potentiation fueled by DRD1-mediated Gαs-βγ dissociation^22,23^ (**Fig. 5C**, 3rd plot). Finally, AC6 alone also generated responses comparable to wild-type cells with lower background activity than AC7 (**Fig. 5C**, 2nd plot). In general, the responses in all AC conditions could be fit to the operational model of dopamine signaling (**Fig. 5C**, solid lines). Together, these results show that the three AC isoforms analyzed here have distinct DRD1 response profiles, all consistent with known features of adenylyl cyclases.

We next expanded this analysis to include the DRD2-Gαi inhibitory axis. We used forskolin stimulation of ACs to enable analysis of DRD2’s inhibitory effects. AC6 is the sole forskolin-responsive isoform in this pool (**Fig. S5)**, and therefore dominated the DRD2-mediated inhibitory response (**Fig. 5D**). The AC6 response mirrored the wild-type response to DRD2, showing that AC6 carries the measured Gαi inhibition (R^2^ = 0.89; **Fig. 5D**). Because AC7 and AC9 are not forskolin-responsive, this assay cannot test their Gαi sensitivity or rank Gαi-responsiveness across isoforms. These measurements provided effective quantitative interaction parameters for AC6.

We next asked whether these individual-AC parameters could predict the behavior of co-expressed AC pairs. The integrated dose responses to pairwise AC combinations could be explained as expression-weighted sums of the single AC response curves, with no additional interaction terms required (R^2^ = 0.94 to 0.97; **Fig. 5E**).

Finally, we challenged the empirically parameterized Isoform-resolved AC Integration Model to predict the integrated cAMP landscape of cells co-expressing both DRD1 and DRD2. Locking the previously derived single-isoform parameters, the model successfully captured the complex two-dimensional signaling surfaces at both 100 nM (**Fig. 5F**, upper rows) and 100 µM (lower rows) dopamine across all eight AC contexts (R^2^ = 0.876 across 847 bins).

These data show how AC isoform expression profiles determine the integrated dopamine response. Manipulating the AC pool actively shifted the apparent dopamine sensitivity of the cell. For example, the AC6+AC7 context sensitized the DRD1 response to lower dopamine concentrations compared to either isoform alone. Furthermore, contexts retaining AC7 exhibited a distinct band of synergistic cAMP elevation in high-DRD2 and medium-DRD1 regimes. This behavior is consistent with the known synergistic stimulation of AC7 by Gβγ subunits released during DRD2 engagement^22^. However, definitively isolating the Gβγ contribution will require targeted scavenging perturbations in future studies.

Together, these perturbations demonstrate that complex signal integration between opposing receptors can be explained from AC diversity, without invoking physical receptor crosstalk. Additionally, by treating the combined effector pool as a transcript-weighted sum, one can potentially predict how diverse cell types, given their AC expression profiles, should integrate conflicting dopaminergic inputs. We next explore this predictive principle using native brain transcriptomic data.

### Model-based simulations predict cell-type-specific dopamine integration in striatal neurons

Our in vitro experiments and predictive mathematical framework demonstrate that the cAMP response of the DRD co-expressing cell is dictated by its adenylyl cyclase composition. To test how this integration logic could apply to native neural circuits in silico, we analyzed single-cell transcriptomes of 14.6 million human neurons (CellxGene Census; Methods). In the striatum, dopamine receptor expression partitioned canonically: direct-pathway MSNs (dMSNs) were DRD1-dominated, indirect-pathway MSNs (iMSNs) were DRD2-dominated, and a rarer population of eccentric MSNs (eMSNs, representing ~5% of clusters) co-expressed both receptors at comparable levels (**Fig. 6A**).

To extrapolate our computational model to these neuronal populations, we examined their AC expression profiles. All three MSN subtypes shared a conserved effector core dominated by the dual-regulated integrator AC5 and the weak Gαs activator AC9. However, their variable AC pools diverged (**Fig. 6A**). eMSNs upregulated AC2 by an order of magnitude compared to dMSNs and iMSNs, while downregulating AC1.

Crucially, the AC expression profiles of these native neurons functionally mirror the components we characterized experimentally in CHO cells (**Fig. 6B**). In MSNs, the primary integrator is AC5, which is biochemically analogous to the CHO cell’s AC6, as both are stimulated by Gαs and inhibited by Gαi. Similarly, the eMSN-enriched AC2 has been shown, like AC7, to act as a conditional amplifier that is insensitive to Gαi but potentiated by Gβγ when Gαs is active^22,23^. This makes it functionally homologous to the AC7 isoform we parameterized in vitro. Because of this conserved biochemical logic, we extrapolated our mathematical framework to computationally simulate how distinct AC profiles could govern signal integration in vivo.

We first simulated the single-receptor-dominated dMSNs and iMSNs under dopamine-only conditions (**Fig. 6C**). In these canonical populations, the hypothetical introduction of eMSN-specific AC2 into the model did not alter the simulated dopamine profiles. Because AC2 requires concurrent Gαs and Gαi/Gβγ drive to act as an amplifier, our model predicts that AC2 is dispensable in cells strictly expressing a single receptor subtype..

In simulations of eMSNs, however, the presence of AC2 dictated the cell’s responsiveness (**Fig. 6D**). Because eMSNs co-express DRD1 and DRD2, a single dopamine transient inevitably drives conflicting Gαs and Gαi inputs. When simulated under dopamine-only conditions, the wild-type (WT) eMSN profile avoids signal cancellation. The conditional amplification of AC2 buffers the predicted response, producing an initial signaling peak before plateauing at a lower, sustained amplitude. If AC2 is reduced to the canonical dMSN level, this buffering is lost, and the simulated cAMP signal collapses to near-zero at saturating dopamine concentrations.

Finally, we extended this principle beyond single-ligand auto-conflict to computationally evaluate how the AC layer resolves the integration of multiple co-transmitted neurotransmitters. Native striatal circuits continuously integrate diverse signals. A classic, well-described example of this is the opposing Gαi drive from dopamine and Gαs drive from adenosine in iMSNs^24^. By verifying the expression of the Gαs-coupled adenosine receptor ADORA2A within our transcriptomic data (**Fig. 6A**), we simulated concurrent dopamine and adenosine receptor activation to evaluate multi-ligand co-stimulation (**Fig. 6E**). Under these conditions, the cAMP response in ADORA2A-expressing iMSNs remained heavily suppressed near baseline across all simulated dopamine concentrations. This reflects classical AC5-mediated Gαi suppression, where DRD2 activation effectively neutralizes the concurrent Gαs drive from adenosine

However, applying this co-stimulation framework to the eMSN population reveals a different integration logic. If AC2 is reduced to the canonical dMSN level, the eMSNs do not produce cAMP under dual stimulation. In contrast, the wild-type eMSN that expresses AC2 amplifies its response through the Gβγ-potentiated amplifier AC2. Consequently, while dMSNs and eMSNs exhibit distinct output profiles in response to dopamine alone, concurrent adenosine stimulation results in a similar cAMP response in both eMSNs and dMSNs.

While our operational model does not explicitly simulate AC1 (as it was absent from our in vitro parameterization), the transcriptomic data suggests its omission does not compromise our predictions. In single-receptor dMSNs and iMSNs, AC1 is likely of lesser importance because these cells lack the opposing G-protein drive necessary to trigger cross-pathway interference. In contrast, our framework predicts that expressing AC1 would be actively detrimental in eMSNs. Because AC1 is attenuated by Gβγ, the concurrent activation of D1 and D2 receptors in eMSNs would generate high levels of free Gβγ that effectively shut down AC1 signaling. By selectively downregulating AC1, eMSNs appear to avoid this auto-conflict. We propose that by minimizing the Gβγ-attenuated AC1 while simultaneously upregulating the Gβγ-potentiated AC2, eMSNs possess a variable AC pool structurally optimized to sustain and amplify signaling during multi-receptor activation.

## Discussion

The prevailing mode of dopamine signaling assumes D1-like and D2-like receptors function independently in separate neuronal populations. However, it has remained unclear how neurons co-expressing those opposing receptors reconcile concurrent Gαs and Gαi signals from the same ligand. Our CHO cell experiments demonstrate that the cAMP output is a weighted sum of single-receptor responses filtered through the AC layer. Thus, while non-additive GPCR effects are often attributed to receptor cooperativity^21^, our results show that substantial signal processing occurs downstream, dictated by the specific adenylyl cyclase (AC) expression profile and its differential sensitivities to Gαs, Gαi, and Gβγ.

Previous studies established ACs, particularly AC5, as integration nodes. However, these studies predominantly focused on how ACs resolve multi-ligand co-stimulation of unrelated receptors, such as integrating a dopamine peak against an acetylcholine dip^25^, or mediating classical competition between co-transmitted dopamine and adenosine^24^. During co-stimulation, the opposing inputs can vary independently in timing, duration, and magnitude. Dopamine receptor co-expression presents a distinct integration mode. Because DRD1 and DRD2 frequently co-localize in unpartitioned spaces, such as *Drosophila* presynaptic active zones^26^and mammalian dendritic spines^9^, both receptors are likely to sense the same dopamine transients and drive concurrent Gαs and Gαi signaling.

In the context of concurrent signaling, modifying the AC composition in CHO cells reshapes the signaling landscape: relying on the Gαi-sensitive AC6 results in strict signal cancellation, whereas introducing the Gαi-insensitive AC9 increases sensitivity to low dopamine. Further incorporating the Gβγ-potentiated AC7 overrides DRD2-mediated inhibition. Thus, by upregulating specific AC isoforms, the cell generates non-additive interactions that receptor tuning alone cannot achieve.

These non-additive interactions may support specific biological functions in native neural circuits. As suggested by our transcriptomic analysis and simulations, the rare population of striatal eccentric MSNs (eMSNs) upregulates the conditional amplifier AC2, the biochemical homolog to AC7 (**Fig. 6A**). We hypothesize that this specific effector profile is used to respond to concurrent DRD1 and DRD2 activation. In the model, omitting this effector switch causes concurrent receptor activation across increasing dopamine concentration, resulting in near-complete signal cancellation (**Fig. 6C,D**). With AC2, the eMSN cells can maintain a sustained, albeit lower, cAMP amplitude at saturating dopamine levels (**Fig. 6D**). Furthermore, similar signal processing behavior could extend to adenosine, whose receptors are co-expressed with dopamine receptors in iMSN cells (**Fig. 6E**). If dopamine and adenosine are simultaneously present, the AC2 layer could respond to Gαs and Gβγ drives to synergistically amplify cAMP responses.

The key features of receptor co-expression and variable AC expression profiles occur in non-mammalian systems (**Fig. S9)**. For example, in *Drosophila* reward (PAM) neurons, D1-like and D2-like receptors co-localize at presynaptic active zones, amplifying low-intensity dopamine signals while clipping high-intensity ones. Although this modulation relies likely on differing receptor affinities, both receptors ultimately converge on a shared cAMP pool. Our transcriptomic analysis reveals that PAM neurons and downstream Kenyon cells co-express the same dopamine receptor pairs while expressing different AC profiles. These comparative data align with our framework, in which neuronal types could differently resolve shared dopamine inputs through their unique AC pools.

In primary MSNs, AC5 is concentrated at the plasma membrane while AC9 localizes to endosomes^27^. Because membrane-localized AC5 is heavily Gαi-inhibited, the local cAMP pool at the cell surface may be dominated by DRD2-mediated inhibition. Conversely, because AC9 is Gαi-insensitive, our framework predicts that endosomal cAMP could effectively evade DRD2-mediated inhibition. This could allow the endosome to maintain a robust, D1-like stimulatory response to drive cAMP production, despite the simultaneous activation of inhibitory receptors at the cell surface.

Several limitations of this study could provoke further investigation of dopamine signaling. First, CHO cells allow high-throughput analysis of the pathway but are far from the neural context where it is most relevant. It will therefore be important to interrogate native neurons with specific endogenous AC profiles. Second, temporal dynamics dopamine could influence integration. Native dopamine signals often occur as rapid phasic bursts and cAMP response is shaped by the dynamic interplay between adenylyl cyclases and cAMP-degrading phosphodiesterases^28^.

Systematically mapping the kinetic space of divergent receptor and AC activation will require high-resolution, time-resolved cAMP measurements. Third, resolving whether concurrent signals are processed differently at the plasma membrane versus endosomes as mentioned above requires compartment-specific cAMP biosensors^29,30^. Finally, it remains unclear whether Gβγ potentiation drives synergistic amplification in native circuits. Confirming the role of this pathway requires targeted Gβγ-scavenging assays.

Our experimental and transcriptomic data suggest a broader conceptual framework: neuromodulatory GPCR pathways can be viewed as flexible computational input-output systems that are structurally analogous to multi-layer neural networks^31^ (**Fig. 7**). In this architecture, a ligand input is processed by a “hidden layer” of surface receptors and G-protein transducers, where cell-type-specific transcript abundances determine the connection weights of each signaling branch. The adenylyl cyclase pool forms the final non-linear integration layer, acting similarly to mathematical activation functions. Just as activation functions dictate how a computational node processes inputs, the biochemical regulatory logic of each expressed AC isoform defines the non-linear transfer function for that branch, utilizing integrators like AC5 and AC6 for standard sigmoidal activation or inhibition, conditional amplifiers like AC2 and AC7 for Gβγ-driven potentiation, or Gαi-insensitive isoforms like AC9 to increase dopamine sensitivity.

**Figure 7.**
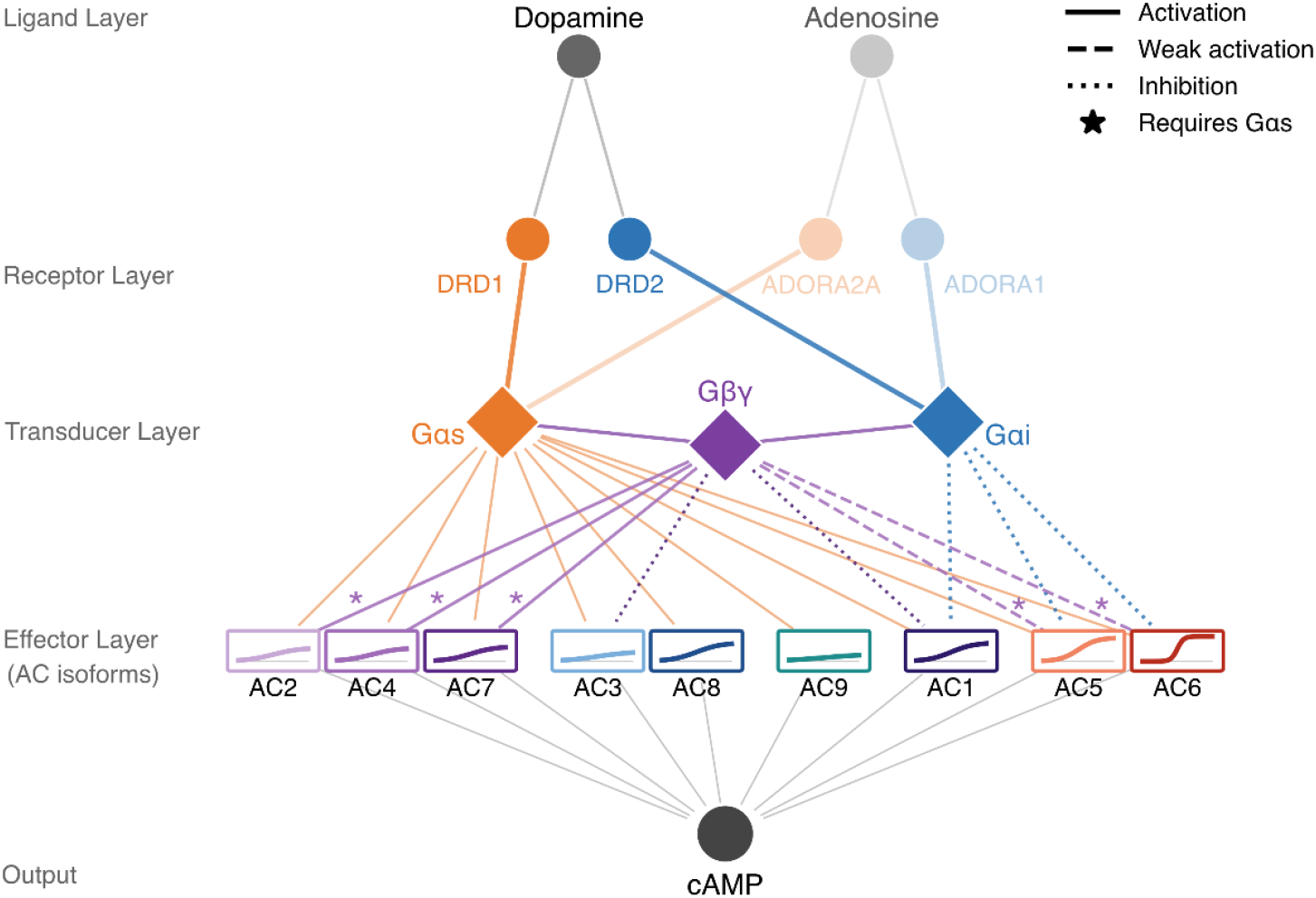
A neural network-like architecture for neuromodulatory GPCR signal integration. Extracellular ligands serve as inputs that are processed through a layer of surface receptors and G-protein transducers, where cell-type-specific transcript abundances act akin to connection weights. The adenylyl cyclase (AC) pool serves as the final non-linear integration layer. In this conceptual framework, the biochemical regulatory logic of each expressed isoform functions similarly to an activation function, enabling specific enzymes to uniquely amplify, integrate, or isolate incoming signals based on their distinct G-protein sensitivities. By varying expression levels across these receptor, transducer, and effector layers, native cell types can rewire their internal processing logic to compute diverse outcomes, such as cancelling, scaling, or comparing conflicting co-transmitted inputs, prior to the final cAMP output.

Viewed through this lens, distinct native cell types represent differently trained biological networks. By varying transcript profiles across receptor and effector layers, a cell type can dynamically rewire its internal processing rules to amplify, cancel, or compare conflicting inputs prior to the final cAMP output. This multi-layer architecture likely represents a generalized computational strategy across the nervous system. Antagonistic receptor pairs are a recurring feature across diverse neuromodulatory networks, including serotonin, adenosine, and histamine systems. In each of these pathways, the rules governing co-activation are not fixed properties of the surface receptors themselves, but are computed by the specific adenylyl cyclase isoform combinations that receive them. Ultimately, a cell type’s local pathway expression profile hardwires its unique computational signature, determining how identical extracellular inputs are transformed into distinct physiological outputs^32,33^.

## Author contributions

J.G. conceived, designed, and performed experiments; analyzed experimental data; developed and fit mathematical models; and performed bioinformatic analysis. M.B.E contributed to experimental design and analysis. J.G. and M.B.E. wrote the paper.

## Acknowledgements

J.G. was supported by the Boehringer Ingelheim Fonds PhD fellowship. M.B.E. is a Howard Hughes Medical Institute Investigator.

## Competing Interests

M.B.E. is a scientific advisory board member or consultant at TeraCyte, Plasmidsaurus, and Spatial Genomics.

## Methods

### Materials Availability

The reagents, ligands, siRNAs, and qPCR probes used in this paper are described in Tables 1 (siRNA sequences) and 2 (qPCR primers), and can be purchased from commercial sources. Plasmids and cell lines generated for this paper, including the dopamine receptor expression constructs of Table 3, are available from the corresponding author upon request.

**Table 1:**
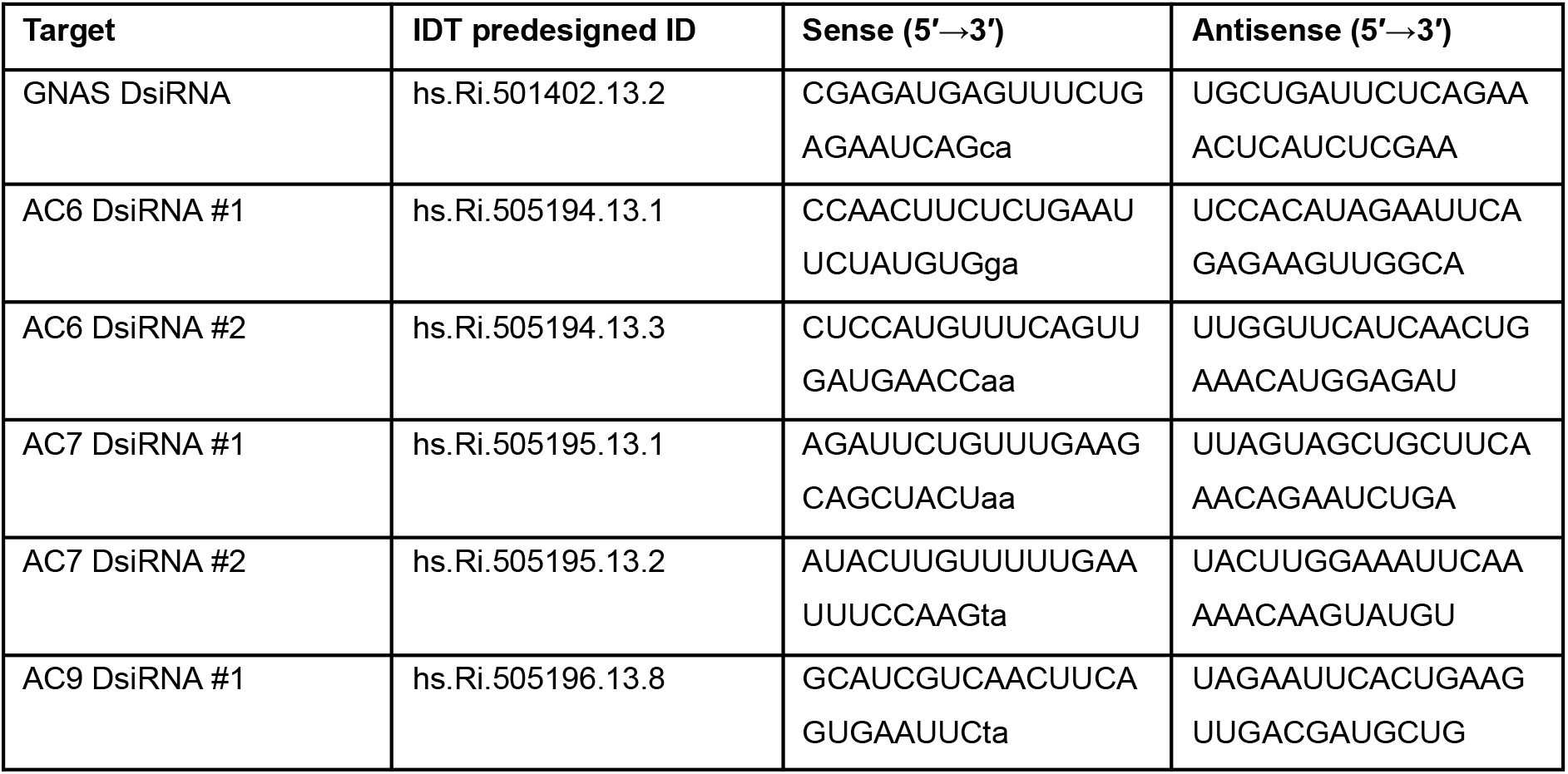
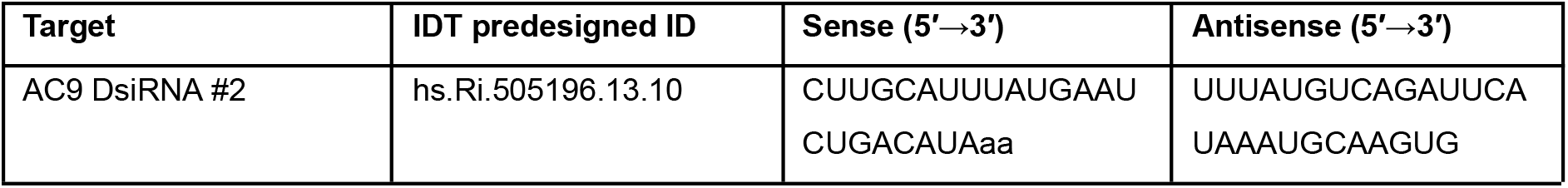
siRNA sequences (Dicer-substrate DsiRNA duplexes; all from Integrated DNA Technologies (IDT)). Sense and antisense strands shown 5′→3′; lowercase residues at the sense 3′ end are DNA overhangs per the IDT 25/27mer DsiRNA design.

**Table 2:**
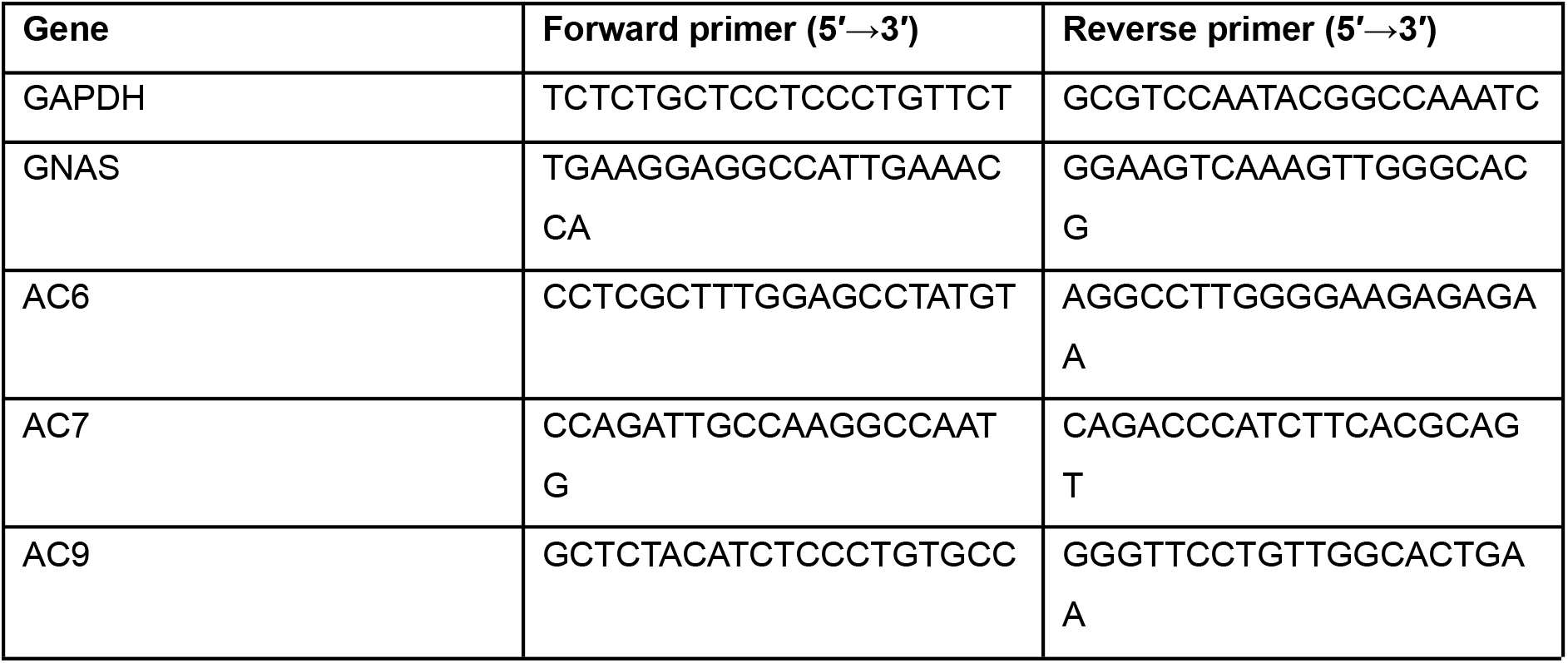
qPCR primers (all from Integrated DNA Technologies (IDT)). Used with SsoAdvanced Universal SYBR Green Supermix (Bio-Rad 1725271) on a Bio-Rad CFX96 Touch.

**Table 3:** Receptor expression constructs. N-terminally tagged dopamine-receptor mRNA expression constructs used in this study. All inserts were synthesized by Twist Bioscience as codon-optimized fragments and cloned downstream of an N-terminal influenza HA signal peptide and a HaloTag7 (HALO) or SNAPf (SNAP) tag, separated from the receptor coding sequence by a GGSGGS linker, in a T7-driven mRNA template plasmid (see Methods, “Dopamine receptor expression plasmids and mRNA production”). Tag versions were verified by sequence analysis: HaloTag7 was identified by the diagnostic A172T mutation^35^ in the absence of the Q165H/P174R double mutation that defines HaloTag9; SNAPf was identified by the E30R substitution that distinguishes the fast-labeling variant from the original SNAP-tag^36^. DRD4 is expressed as the 4-repeat (4R) allele of the exon-3 VNTR, not the 7-repeat allele.

| Construct | Receptor (variant, length) | RefSeq mRNA | Tag (N-terminal) | Tag reference |
| --- | --- | --- | --- | --- |
| HA-HaloTag7-DRD1 | Human DRD1 (canonical, 446 aa) | NM_000794 / NP_000785 | HaloTag7 | Encell et al. 2012 |
| HA-HaloTag7-DRD2 | Human DRD2 (long isoform, 443 aa) | NM_000795 / NP_000786 | HaloTag7 | Encell et al. 2012 |
| HA-SNAPf-DRD2 | Human DRD2 (long isoform, 443 aa) | NM_000795 / NP_000786 | SNAPf | Sun et al. 2011 |
| HA-HaloTag7-DRD3 | Human DRD3 (canonical, 400 aa) | NM_000796 / NP_000787 | HaloTag7 | Encell et al. 2012 |
| HA-HaloTag7-DRD4 | Human DRD4 (4-repeat allele, 419 aa) | NM_000797 / NP_000788 | HaloTag7 | Encell et al. 2012 |
| HA-HaloTag7-DRD5 | Human DRD5 (canonical, 477 aa) | NM_000798 / NP_000789 | HaloTag7 | Encell et al. 2012 |

### Cell lines and culture conditions

All experiments were performed using the Chinese hamster ovary K1 subline (CHO-K1), obtained from ATCC (Cat# CCL-61; female, adult ovary origin). This cell line was selected because it lacks endogenous dopamine receptor expression while retaining the core downstream transduction machinery relevant to this study: Gαs (*GNAS*), three Gαi isoforms (Gαi1, Gαi2, and Gαi3), and three adenylyl cyclase isoforms (AC6, AC7, and AC9). Crucially, CHO-K1 does not express the Gαo subunit, providing a clean background to explicitly test Gαo-dependence for receptors such as DRD3 and DRD4. The absence of endogenous dopamine receptors (*DRD1–DRD5*), as well as the expression profiles of the relevant transducers and adenylyl cyclases, were confirmed via bulk RNA-seq of the parental CHO-K1 line (Figure S1).

Cells were cultured in Alpha-MEM (Gibco, Cat# 12561056) supplemented with 10% fetal bovine serum (VWR, Cat# 311K18), 1 mM sodium pyruvate (Gibco, Cat# 11360070), 2 mM L-glutamine (Gibco, Cat# 25030081), 1 U/mL penicillin, and 1 μg/mL streptomycin. Cells were maintained at 37°C under 5% CO2 in a humidified incubator. Cultures were passaged at 90% confluence using StemPro Accutase Cell Dissociation Reagent (Gibco, Cat# A1110501) and used between passages 4 and 25 post-thaw.

### Reporter cell line construction (CHO-K1::cAMPinG1)

The stable reporter line utilized in all experiments is a monoclonal CHO-K1 derivative expressing the excitation-ratiometric cAMP biosensor cAMPinG1^34^. cAMPinG1 is a circularly permuted Citrine–PKA-regulatory-domain fusion; upon cAMP binding, it undergoes a conformational change that shifts the relative excitation efficiency between two maxima (~488 nm and ~405 nm). Because the readout is an excitation ratio rather than raw fluorescence intensity, it remains largely insensitive to per-cell expression variability and instrument-level gain settings. This ratiometric stability is essential for the single-cell dose-response binning employed in this study. A Kusabira Orange 525 (KO525) excitation channel provides the ratiometric denominator. At saturating cAMP levels, Citrine exhibits an approximate 1000% ΔF/F and KO525 an approximate 61% ΔF/F. On our instrument (Beckman CytoFlex S), the Citrine/KO525 ratio ceiling at saturating forskolin (20 μM) corresponds to Cit/KO ≈ 20.7.

The cAMPinG1 sensor^34^ was obtained from the RIKEN BioResource Research Center DNA Bank (full list of cAMPinG1 RDB plasmid accessions reported in the supplementary material of Yokoyama et al., 2024) and synthesized as a gene fragment (Twist Bioscience). This fragment was cloned between PiggyBac inverted terminal repeats downstream of a strong constitutive promoter (EF1α), utilizing a puromycin resistance gene as a selection marker. The resulting transfer plasmid was co-transfected with a hyperactive PiggyBac transposase (System Biosciences, Cat# PB210PA-1) into the parental CHO-K1 line using Lipofectamine 2000 (Invitrogen, Cat# 11668019) according to the manufacturer’s protocol. Stable integrants were selected in 10 μg/mL puromycin (Gibco, Cat# A1113803) for 10 days, expanded, and subjected to limiting dilution in 96-well plates to isolate single clones. Candidate clones were expanded and screened for (i) high baseline KO525 signal with low variance, and (ii) a robust Citrine/KO525 ratio increase following acute stimulation with 10 μM forskolin (MedChemExpress, HY-15371). The final validated monoclonal line, CHO-K1::cAMPinG1, was used for all reported reporter assays.

### Dopamine receptor expression plasmids and mRNA production

Human DRD1 (RefSeq NM_000794), DRD2 long isoform (NM_000795), DRD3 (NM_000796), DRD4 4-repeat variant (NM_000797), and DRD5 (NM_000798) coding sequences were synthesized by Twist Bioscience, codon-optimized for mammalian expression, and cloned downstream of an N-terminal HaloTag7^35^ or SNAPf^36^ tag with a short flexible linker (GGSGGS) in a T7-driven mRNA template plasmid. The full N-terminal architecture of the expressed protein is: ATG → influenza hemagglutinin (HA) signal peptide (16 aa, MKTIIALSYIFCLVFA, included to ensure cotranslational ER targeting and surface trafficking) → HaloTag7 or SNAPf → GGSGGS linker → receptor coding sequence. All five receptors were cloned in HaloTag7-tagged form for the single-receptor experiments of Figures 3A and S3; DRD2 was additionally cloned in SNAPf-tagged form for the orthogonal two-color combinatorial DRD1+DRD2 experiments of Figures 2E, 4, and 5. Construct details are summarized in Table 3. Linear in vitro transcription templates were generated by PCR amplification from each plasmid using a 5’ primer and a 3’ Ultramer primer (IDT), the latter encoding a templated 120-nt polyA tail. PCR products were column-purified before use as IVT templates. In vitro transcription was carried out with T7 RNA polymerase (NEB M0251), an NTP mix of ATP, GTP, and CTP (NEB), Pseudouridine-5’-triphosphate (TriLink N-1019) in place of UTP to increase mRNA stability, and CleanCap Reagent AG (TriLink N-7113) for co-transcriptional 5’-capping. mRNA was purified using RNA Clean & Concentrator-25 (Zymo Research, R1018), resuspended in nuclease-free water, quantified by NanoDrop, and stored at −80 °C in single-use aliquots.

### Transfection protocols

CHO-K1::cAMPinG1 cells were seeded into 12-well plates at 1.5 × 10^5 cells/well 24 h before transfection. For mRNA-based expression of dopamine receptors, cells were transfected with Lipofectamine MessengerMAX (Invitrogen LMRNA003) following the manufacturer’s mRNA protocol, with per-well mRNA amounts titrated to yield the intended surface-abundance distribution (50–500 ng per well across different conditions). For siRNA knockdown, unless specified, Lipofectamine RNAiMAX (Invitrogen 13778-100) was used at 33 nM final siRNA (IDT), delivered by reverse transfection at seeding (full schedule below); receptor mRNA was added 24 h later.

### siRNA knockdown for GNAS and adenylyl cyclase isoforms

Pooled siRNAs (IDT) for adenylyl cyclase isoforms and GNAS were used throughout to reduce off-target risk. Each pool comprises four independent oligonucleotides targeting distinct sites in the target mRNA (sequences in Table 1). For single-knockdown experiments, cells were reverse-transfected with 10 nM siRNA at the time of seeding using Lipofectamine RNAiMAX, with receptor mRNA co-transfection performed at 24 h. For combined knockdowns (siAC6+siAC7, siAC7+siAC9, siAC6+siAC7+siAC9), siRNAs were pooled to a combined final concentration of 10 nM, with each component present at 5 nM; equivalent siNT was added to single-knockdown wells to match total siRNA concentration across conditions.

### Surface labelling with SNAP- and HALO-Surface ligands

At 30 min before flow cytometry acquisition, cells were labelled with cell-impermeable SNAP-Surface and/or HALO-Surface ligands to report surface receptor abundance. Single-colour experiments used SNAP-Surface® Alexa Fluor® 647 (NEB S9136S) at 1 μM for SNAP-tagged receptors or Janelia Fluor® 549i HaloTag® Ligand (Promega HT1080) at 100 nM for HALO-tagged receptors. Combinatorial DRD1 and DRD2 experiments combined both ligands at the same per-ligand concentrations in a single labelling step. Labelling was performed in complete growth medium at 37 °C for 30 min, followed by three washes with warm complete medium to remove unbound ligand.

### Dopamine stimulation and flow cytometry acquisition

Cells were dissociated gently with Accutase to preserve the surface receptors and tags. The Accutase was neutralized with flow cytometry buffer (HBSS containing 2.5 mg/mL Bovine Serum Albumin (BSA)) enriched with 100 μM IBMX (MedChemExpress) to inhibit phosphodiesterases and prevent cAMP degradation. Cells were then filtered through a 40 μm mesh into 96-well plate and treated with dopamine (MedChemExpress, HY-B0451A) at concentrations spanning 1 nM to 100 μM. For experiments involving Gαi-coupled receptors (DRD2, DRD3, DRD4), adenylyl cyclase was activated with 10 μM forskolin (MedChemExpress, HY-15371) 5 minutes post-dopamine treatment. The Citrine/KO525 excitation-ratio reaches steady state and stays stable after 5 minutes (continuous-flow kinetics experiments, Fig. S2) and flow cytometry analysis was performed within this time window.

Acquisition was performed on a Beckman CytoFlex S flow cytometer equipped with 405, 488, and 638 nm excitation lasers. cAMPinG1 Citrine was acquired with 488 nm excitation and a 525/40 BP filter, and KO525 with 405 nm excitation and a 585/42 BP filter; the Citrine/KO525 excitation ratio (Cit/KO) is the primary cAMP readout. HALO-tagged receptor labelled with Janelia Fluor® 549i was acquired with 488 nm excitation and a 585/42 BP filter, and SNAP-tagged receptor labelled with Alexa Fluor® 647 with 638 nm excitation and a 660/10 BP filter.

### RT-qPCR for knockdown validation and AC isoform profiling

Total RNA was isolated 48 h after siRNA knockdown using TRI Reagent (Zymo Research, R2050-1-50) and Direct-Zol RNA Miniprep Kit (Zymo Research, R2050). RNA concentration and purity were measured by NanoDrop. cDNA was synthesized from 300 ng total RNA using iScript Reverse Transcription Supermix (Bio-Rad 1708840) in 20 μL reactions. qPCR was performed on a Bio-Rad CFX96 Touch using SsoAdvanced Universal SYBR Green Supermix (Bio-Rad 1725271) in 10 μL reactions with 300 nM each primer and 2 μL of 1:5 diluted cDNA. Primer sequences for GNAS, AC6, AC7, AC9, and the housekeeping gene GAPDH are given in Table 2. Expression was quantified by the ΔΔCq method against GAPDH, and knockdown efficiency was computed as 1 − 2^(−ΔΔCq), i.e., one minus the fold change of the target gene in the siTarget condition relative to siNT (i.e., 1 − (target in siTarget) / (target in siNT)). Across four independent RNA isolations, the consensus knockdown efficiencies were: siGNAS 92–97%, siAC6 (pool) 65–83%, siAC7 (pool) 53–72%, siAC9 (pool) 69–86%, and siTriple (AC6+AC7+AC9) 60–78% per isoform. AC isoform fractions in CHO-K1 were computed from the pooled WT + siGNAS + siNT samples (n = 7 independent isolations) as AC6: 65.6 ± 10.0%, AC7: 11.3 ± 2.2%, AC9: 23.1 ± 8.1% of the sum (AC6 + AC7 + AC9) on a ΔCq-converted relative-abundance scale; these qPCR-derived fractions were used in the three-AC layer model of Figure 4.

Analyses were implemented in Python; custom code is available in the repository described in the Methods, organized by figure panel. Substantial portions of the analysis code were developed with the assistance of Claude (Anthropic); the author retains full responsibility for the scientific content and conclusions. The subsections below describe each analytical step: external data curation (dopamine binding-affinity curation), single-cell transcriptomic analysis of striatal medium spiny neurons, flow cytometry data processing (gating, binning, normalization, contamination correction, and sensor-saturation truncation), parameter fitting and inference (per-receptor and multi-condition operational models, model comparison, transducer-level inference, and batch-matching), and cell-type-specific signaling simulations.

### Dopamine binding affinity curation

Equilibrium dissociation constants (Ki) for dopamine at human DRD1–DRD5 were curated from the IUPHAR/BPS Guide to Pharmacology (GtoPdb; guidetopharmacology.org). We queried for dopamine as the ligand across all five receptor subtypes and applied a stringent filter: only human recombinant receptor assays using radioligand competition binding (reported as pKi or pKd) were retained. Assays using non-human species, tissue preparations, or functional readouts were excluded.

The curated dataset comprises 24 individual measurements: DRD1 (N=6), DRD2 (N=5), DRD3 (N=10), DRD4 (N=2), and DRD5 (N=1). Values are presented as pKi (= −log_10_ Ki), with median Ki in nanomolar annotated per subtype. The data reveal approximately three orders of magnitude variation in dopamine affinity across subtypes: DRD3 (median 20 nM) and DRD4 (8 nM) are high-affinity; DRD2 (257 nM) and DRD5 (229 nM) are intermediate; and DRD1 (2,356 nM) is the lowest-affinity subtype.

The full unfiltered distribution of retrieved affinities and the per-criterion exclusion breakdown are shown in Fig. S6.

### Single-cell RNA-seq data acquisition and per-cell extraction

Single-cell gene expression data were obtained from the CellxGene Census (Chan Zuckerberg Initiative; cellxgene.cziscience.com), release 2025-01-30, accessed through the cellxgene-census Python API. For each Census dataset, a server-side query selected human (Homo sapiens) primary-data cells annotated as brain tissue and retrieved the raw UMI counts for a 28-gene panel (the five dopamine receptors DRD1–DRD5, all ten adenylyl cyclases ADCY1–ADCY10, four adenosine receptors ADORA1, ADORA2A, ADORA2B and ADORA3, the PDE4 phosphodiesterases PDE4A–D, and five Gα subunits GNAS, GNAI1–GNAI3 and GNAO1), together with each cell’s library size (total UMI), cell-type annotation, tissue and anatomical-region (UBERON) labels, assay, donor, and dataset identifiers. Cells were then restricted to neuronal cell types by matching the CellxGene cell-type annotation. Per-dataset extractions were merged into a single per-cell table; data quality was assessed by per-cell-type detection-rate summaries and a positive control confirming the expected co-detection of ADORA2A and DRD2 in striatal neurons. Raw UMI counts were used throughout; no normalization, log-transformation, or imputation was applied. Genes were scored as detected at a threshold of one or more UMI, and cells were used as deposited in the Census (author-level quality control; primary-data cells only). Because the key analyses depend on a small co-expressing population, we confirmed that the result is not driven by low-quality cells: imposing standard per-cell quality filters (a minimum of 1,000 total UMI and 500 detected genes per cell) removed essentially no striatal medium spiny neurons (1 of 12,150) and left the DRD1/DRD2 co-expression fractions unchanged. Extractions were run in parallel across datasets; all pipeline code and the dataset manifest are provided in the deposit.

### Striatal MSN expression profiles

From the per-cell table, the dopamine receptors (DRD1, DRD2), core adenylyl cyclases (ADCY5, ADCY9), variable adenylyl cyclases (ADCY1, ADCY2), and adenosine receptors (ADORA1, ADORA2A) were quantified in direct-pathway (dMSN), indirect-pathway (iMSN), and eccentric medium spiny neurons (eMSNs), the last defined as striatal medium spiny neurons carrying the atlas “medium spiny neuron” annotation without a direct- or indirect-pathway assignment; consistent with the eccentric MSNs of ref. 10, these cells co-express DRD1 and DRD2 (Fig. 6A), separately for the caudate nucleus and putamen. Per-cell expression was computed as counts per million (CPM = gene UMI / total UMI × 10^6^) and averaged within each cell-type × region group (groups with fewer than 50 cells were excluded). The resulting per-group means are the stacked expression profiles shown in Fig. 6A.

### Cell-type-specific signaling simulations

To project the integration model onto native neurons, the operational AC-integration model (Appendix 1) was evaluated using each MSN subtype’s adenylyl cyclase composition. The biochemical parameters were taken from the in-vitro fits: per-isoform transducer parameters from the joint per-AC fit (Fig. 5C), DRD1 and DRD2 receptor parameters from the single-receptor fits (Fig. 3), the Gαi-inhibition coefficients from the DRD2 fit (Fig. 5D), and the cAMPinG1 sensor calibration. Absolute receptor drive levels are not constrained by the transcriptomic data and were set to representative values (surface levels for DRD1 and DRD2, and, for co-stimulation, ADORA2A and ADORA1; DRD2:DRD1 ratios of 1.0 and 1.3); the qualitative eMSN result is unchanged across a range of these levels and ratios (Fig. S10). Dopamine drives DRD1 (Gαs) and DRD2 (Gαi); adenosine drives ADORA2A (Gαs) and ADORA1 (Gαi). Adenosine was applied at a saturating concentration (10 µM), so the co-stimulation result is insensitive to the exact receptor affinities, which were set to representative literature values (adenosine Ki ≈ 10 nM at ADORA1 and ≈ 150 nM at ADORA2A, within the range compiled by Fredholm et al., 2001)^37^. Because the striatal isoforms are biochemically analogous to those characterized in CHO-K1, AC5 was modeled with the AC6 transfer function (Gαs-stimulated, Gαi-inhibited) and AC2 with the AC7 transfer function (Gαs-stimulated, partially Gαi-insensitive). AC2^low^ denotes AC2 reduced to the canonical dMSN level, not its complete removal. Simulations cover the single-receptor-dominated dMSN and iMSN responses (Fig. 6C), the eccentric-MSN response at the eMSN and dMSN AC2 levels (Fig. 6D), and dopamine/adenosine co-stimulation (Fig. 6E).

### Drosophila single-cell analysis

Adult-head single-cell expression was taken from the Fly Cell Atlas^38^, 10x dataset (head; 100,527 cells, 13,056 genes, raw counts). Raw counts were normalized per cell to 10,000 total counts and log1p-transformed. Within each annotated cell type, cells co-expressing both Dop1R1 (D1-like) and Dop2R (D2-like) (raw count > 0 for both) were identified, and for the five adenylyl cyclases (rut, Ac78C, Ac3, Ac13E, Ac76E) we computed, among these co-expressing cells, the percentage expressing each gene and the mean log-normalized expression (Fig. S9). The complete split by co-expression group (Both / Dop1R1-only / Dop2R-only / Neither) is provided with the deposited data. Extraction and plotting code is provided in the deposit.

### Flow cytometry gating and per-cell filtering

Raw flow cytometry files were gated in two stages. Cell-population gating was applied in a custom Python GUI^39^ and used a fixed hierarchy: FSC-A × SSC-A to exclude debris and very large aggregates, followed by a reporter-expressing gate defined as the central 99% of the KO525 fluorescence distribution observed in parental CHO-K1::cAMPinG1 cells. Gated cells were exported as numpy arrays (.npy). Surface-receptor gating on the SNAP- or HALO-surface-label channel (single channel for single-receptor experiments, both channels for combinatorial experiments) was then applied post-hoc in our custom downstream analysis pipeline, with thresholds set once per cell-line batch using the parental line as the SNAP/HALO-negative reference (a per-session unstained control was not collected). An example of this gating strategy is shown in Figure S11.

### Single-cell binning into receptor-expression dose surfaces

For single-receptor experiments, gated cells were binned by surface-label intensity into 8–10 log-spaced bins spanning the receptor-expressing population. For each (bin, dopamine) combination, the per-cell Cit/KO ratio distribution was summarized by its geometric mean (reporter response) with 95% confidence interval computed by cell-level empirical bootstrap (n = 200 resampling iterations, described in Methods). Bins with fewer than 200 cells were pooled with the adjacent bin. For combinatorial (DRD1+DRD2) experiments, gated cells were binned on a two-dimensional grid of HALO-DRD1 × SNAP-DRD2 surface-label intensities using log-spaced bin edges per axis, with the same minimum-cell-count rule applied on a per-grid-cell basis. The resulting measurement is a four-dimensional array: (DA concentration, [D1] bin, [D2] bin, sample) → Cit/KO + bootstrap CI.

### Normalization of the cAMP reporter

The primary reporter readout Cit/KO is anchored on two biological endpoints measured within the same experimental batch. The upper reference (“ceiling”) is the Cit/KO ratio of untransfected CHO-K1::cAMPinG1 cells stimulated with 10 μM forskolin, which saturates the cAMPinG1 sensor (FSK ceiling = 20.74 across batches; per-batch values are given in the analysis metadata). The lower reference (“reporter background”) is the Cit/KO ratio of untransfected, unstimulated CHO-K1::cAMPinG1 (reporter background = 1.03 in the “old” batch, 0.94 in the “new” batch, consistent with a small baseline shift across cell-line batches, possibly reflecting silencing or loss of reporter expression upon longer storage of the frozen stocks). With C the forskolin-saturated ceiling and b the reporter background, the normalized signal is 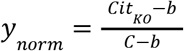, so *y*_*norm*_ = 0 corresponds to the reporter baseline in the absence of Camp elevation and *y*_*norm*_ = 1 corresponds to sensor saturation. For Gαi-coupled receptor experiments the assay operates off a forskolin-elevated baseline; for Gαs-coupled receptors the unstimulated reporter background is the operating point.

### Contamination correction for bleed-through from untransfected cells

Because surface-receptor gating is imperfect at the dim-expression edge, a small fraction of gated cells are in fact untransfected and contribute an untransfected-baseline signal. This contaminating fraction is estimated per bin from the SNAP/HALO intensity distribution of parental (untransfected) control cells. With f the estimated untransfected (contaminating) fraction, the contamination-corrected reporter value is 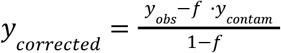. This correction is applied per (bin, dopamine) combination before operational model fitting.

### Sensor-saturation truncation

Near the sensor ceiling, the excitation-ratio readout becomes a nonlinear function of cAMP and small further increases in cAMP no longer produce proportional increases in Cit/KO. To prevent saturated bins from biasing operational-model fits in which the signal is proportional to transducer activation, bins in which the vehicle (no-dopamine) Cit/KO already exceeded 55% of the FSK ceiling were excluded from the fit domain for Gαs-coupled receptors. For Gαi-coupled receptors, which operate off a forskolin-elevated baseline that is intentionally close to saturation, the inhibitory excursion from that baseline was fit instead of the absolute signal, and the truncation did not apply. The 55% cutoff was set conservatively below the sensor’s apparent saturation regime: bins in which the unstimulated baseline already occupies more than half of the dynamic range cannot reliably register further receptor-driven cAMP increase, and including them biases the apparent transducer-pool half-saturation *K*_*m*_ downward.

### Single-receptor operational-model fits

For each receptor, the modified operational model (Appendix Eq. A1.20) is fit to the binned single-cell dose-response data of Figures 3 and S3. The dopamine binding affinity *K*_*d*_ is locked at the median pK_1_ from the IUPHAR/BPS Guide to Pharmacology curated set (Fig. S6 and Methods, “Dopamine binding affinity curation”); the remaining parameters (*n, K*_*m*_, *m, A, y*_0_, ρ_0_) are fit per receptor by non-linear least squares as described under “Parameter fitting” below. Per-receptor fit values with 95% bootstrap confidence intervals are reported in Supplementary Table S1.

Of the five dopamine receptor subtypes assayed in CHO-K1, the operational-model fit is reported for DRD1, DRD2, DRD4, and DRD5; DRD3 is shown as a silent control. DRD3 produces no measurable dose-response in this background because its primary transducer Gαo is not endogenously expressed in CHO-K1. DRD4’s Gαi-coupled response is shallow and has limited dynamic range, consistent with DRD4’s canonical preference for Gαo over Gαi; bootstrap confidence intervals on its fit parameters reach the optimizer bounds, and the corresponding row of Supplementary Table S1 reports point estimates only. DRD5 has a single qualifying entry in the curated affinity dataset; its fit is included in the receptor panel for completeness, with the corresponding profile-likelihood identifiability check shown in Supplementary Figure S7. Quantitative downstream analyses (Figures 3–5) use only DRD1 and DRD2.

### Parameter fitting, model selection, and uncertainty quantification

We used three parameter-fitting strategies, each tailored to the dimensionality and the question that fit was answering.

### Per-receptor operational models

Single-cell flow cytometry data carries high variance and biological outliers. To prevent these outliers from artificially skewing the fit, we fit the per-bin geometric means using a non-linear least-squares algorithm (Trust Region Reflective, implemented in SciPy) paired with a robust soft-L1 loss function rather than the raw χ^2^. Residuals are weighted by per-bin SEM so that better-sampled bins exert proportionally more influence on the fit. Trust Region Reflective natively enforces our physically motivated parameter bounds: amplitude *A* is restricted to positive values for Gαs-coupled receptors (*A*∈[0. 01, 5]) and negative values for Gαi-coupled receptors (*A*∈[− 5, − 0. 01]); transducer-step half-max *K*_*m*_ ∈[1, 10^7^] (in PE-A × occupancy units); transducer-step Hill coefficient *m*∈[1, 5]; binding-step Hill coefficient *n*∈[0. 3, 4]; baseline *y*_0_ ∈[− 0. 5, 0. 5] (Gαs) or [0, 2] (Gαi); basal receptor activity ρ_0_ ∈[0, 0. 5]. The dopamine binding affinity *K*_*d*_ is locked at the median *pK*_*i*_ from the IUPHAR/BPS Guide to Pharmacology curated set (Fig. S6) and not fitted. The same pipeline is used for the DRD1 ± Gαs shared-parameter fit (Figure 3C,D; Table S2) by adding a per-condition transducer-pool variable while keeping the receptor-intrinsic parameters shared.

### Joint multi-condition AC-isoform models

For the multi-condition AC experiments the model must simultaneously explain wild-type and several knockdown contexts: 15 parameters across five isolation conditions for the per-AC DRD1 fit (Figure 5C), and 3 parameters across four conditions for the DRD2 Gαi-inhibition fit (Figure 5D). For these higher-dimensional joint fits we used the gradient-free Nelder-Mead simplex algorithm (implemented in SciPy) to minimize the unweighted sum of squared residuals across all included conditions. Because Nelder-Mead is unconstrained, physical parameter bounds were enforced by returning a severe mathematical penalty whenever the optimizer tested biologically impossible values. An explicit Gβγ-potentiation term for the AC7 isoform was tested and omitted, as it was non-identifiable (degenerate with the isoform’s Gαs-driven amplitude and Gαi-inhibition coefficient) and did not improve the fit (data in the deposit).

### Model-comparison fits

To evaluate alternative structural architectures of the dual-receptor combinatorial integration, we compared three models against the wild-type combinatorial surface (Appendix 1, Models 1–3): the Additive model (no free parameters), the Cooperative model (one free parameter, the heterodimer crosstalk term γ), and the AC Integration model (one free parameter, the Gαs/Gαi competition term κ at AC6). Because these fits are intended for broad model comparison rather than precise parameter estimation, per-bin SEM weighting was omitted (all variants are evaluated against the same error structure, so the relative AIC is unchanged by weight choice) and simpler optimizers were used. For the two single-parameter models the optimum was located with a bounded scalar minimizer (SciPy), preceded by a coarse log-grid scan to bracket the minimum.

### Initialization and robustness

Non-linear models are prone to getting trapped in local minima: suboptimal fits that look correct locally but miss a deeper global optimum elsewhere in parameter space. To find the true global solution, every multi-condition fit was run from many independent starting points. The 15-parameter per-AC DRD1 fit used 41 seeds (one warm-start at the previous best plus 40 random restarts drawn from physically plausible per-parameter ranges using a deterministic RNG seed for reproducibility); the 3-parameter DRD2 fit used 36 seeds sampled from a Cartesian grid over *D*_*FSK*_, ε_6_, and ε_7_. The lowest-cost converged solution across all seeds was selected as the final point estimate.

### Uncertainty quantification

Parameter 95% confidence intervals were computed using two strategies, matched to each fit class. For the per-receptor fits (Supplementary Tables S1–S2), CIs come from a 200-iteration cell-level empirical bootstrap: cells are resampled with replacement within each (date × dopamine concentration) sample, per-bin medians are then recomputed by re-binning the resampled cells on the surface-receptor channel, and the model is refit through the same Trust Region Reflective + soft-L1 pipeline. For the joint multi-condition fits (Supplementary Table S3; Supplementary Fig. S8), CIs come from profile-likelihood scans: each fitted parameter is held fixed on a grid spanning ±1 log decade (log-scaled parameters) or ±0.7 fractional units (linear parameters) around its point estimate, and the remaining parameters are re-fit at each grid point. The 95% interval for each parameter is the range over which Δχ^2^ ≤3. 84 (one degree of freedom; α = 0. 05). Profile-likelihood scans are also performed for the per-receptor fits (Supplementary Figure S7) as a complementary identifiability check; they qualitatively agree with the bootstrap intervals to within a factor of two for the well-fit DRD1 and DRD2 parameters, and document the limited identifiability of DRD4 and DRD5, which are not used in any quantitative downstream analysis.

### Model selection and cross-validation

Model comparisons (single-AC vs three-AC; κ-only vs both-scale on transducer perturbations; κ on WT vs siAC7+9 surfaces; the *K*_*m*_-only vs both-scale fit for the DRD1 ± Gαs comparison) were made by Akaike Information Criterion with finite-sample correction (AICc) and by held-out *R*^2^ on a 20% held-out fraction (cells stratified by bin and by dopamine concentration), re-computed across 10 random held-out splits. AIC and held-out *R*^2^ values cited in the main text are the medians over those splits.

All fitted parameter sets, multi-start logs, per-condition *R*^2^ values, profile-likelihood scans, and bootstrap output are deposited at CaltechDATA, organized by figure panel.

### Inference of transducer level [G] under siRNA knockdown

For experiments involving siRNA-mediated depletion of GNAS, the effective transducer level [G] under knockdown was treated as a single free parameter shared across all receptor-abundance bins within a knockdown condition, while the receptor-intrinsic parameters (n, K□, m, A, y_0_, ρ_0_) were held common to the wild-type and knockdown datasets. Fitting the DRD1 ± siGNAS dose-response surfaces in this way yielded effective transducer levels of [G] = 0.50 of wild-type at 1 nM siRNA and 0.21 at 33 nM (Supplementary Table S2), corresponding to 50% and 79% reductions in the functional Gαs pool. These functional reductions are smaller than the depletion of GNAS transcripts measured by qPCR over the same conditions (91% at 1 nM and 93% at 33 nM; Fig. S4), consistent with a pool of pre-existing Gαs protein that persists at the time of the assay. The fitted [G] values are the ones used to rescale the drive axis in the curve-collapse panel (Fig. 3F).

To test whether the perturbation acts purely through the transducer pool, we compared two models. In the first, only the transducer pool [G] varies per condition (equivalently, a rescaling of the drive/K□ axis), with all receptor-intrinsic parameters shared across wild-type and ΔGs conditions. In the second alternative, the amplitude A is additionally allowed to vary per condition. AICc favored the [G]-only model by ΔAICc = 30.1, indicating that Gαs depletion is consistent with pure transducer depletion (Eq. A1.15) rather than a change in receptor coupling efficiency. Per-condition shared-parameter fit values are reported in Supplementary Table S2.

### Experimental batches and batch-matching

Two experimental batches contributed to the results in this study. The “old” batch (February 6–10, 2025) covered the single-receptor dose-response panels (Figures 3 and S3), with mRNA doses in the low and high ranges, and had an instrument background fluorescence of 5.28 arbitrary units. The “new” batch (February 27, 2025 onwards) covered the transducer-perturbation experiments (Figure 3) and all adenylyl cyclase knockdown experiments (Figures 4 and 5), with low and intermediate mRNA doses and an instrument background fluorescence of 3.41 arbitrary units. Batches are never mixed within a figure panel. The forskolin-defined cAMP ceiling and the reporter background used for normalization were measured per batch from untransfected control cells acquired in the same cytometry session as the experimental samples.

### Statistics and reproducibility

#### Replicates and sample sizes

Replicate numbers are reported per panel. Throughout, a biological replicate is an independent experiment performed on a separate date (separate transfection and flow-cytometry session); within each replicate, every surface-receptor-abundance bin summarizes a large single-cell population (more than 100,000 single cells were acquired per replicate before gating and binning). Single-receptor operational fits used n = 4 independent dates for DRD1 and n = 5 for DRD2 (Fig. 3A,B; Fig. S3). The dual-receptor combinatorial surfaces used n = 3 independent dates (Fig. 4). Adenylyl cyclase knockdown dose-responses used n = 2 to 4 independent dates per AC context (Fig. 5C to 5E). RT-qPCR fold changes used n = 3 to 14 independent qPCR runs per condition-and-gene combination (Fig. 5B; exact values in Supplementary Table S4). Sample sizes were not predetermined by a power analysis; they reflect the number of independent experiments accumulated for each condition.

#### Data representation and error bars

Dose-response points are per-bin medians of the single-cell distribution, plotted against measured surface-receptor abundance, and error bars are ±1 SEM across independent replicate dates. RT-qPCR bars are mean ± SEM across independent runs. In the two-dimensional co-expression heatmaps (Fig. 2E, Fig. 4, Fig. 5F), each tile is the median normalized cAMP of its DRD1-by-DRD2 abundance bin; tiles containing fewer than 50 cells were left unfilled (grey) and excluded from fitting and evaluation.

#### Statistical tests

The only frequentist hypothesis tests in this study are the RT-qPCR knockdown comparisons (Fig. 5B and Fig. S4). For each transcript, the per-run log_2_ fold change relative to non-targeting control was compared against a null of 0 with a one-sample, two-tailed t-test (degrees of freedom = n − 1); exact t, degrees of freedom, and P values for the Fig. 5B panel are given in Supplementary Table S4, and the Gαs-knockdown value (Fig. S4) is reported in that figure legend. In the figures these are abbreviated as *P < 0.05, **P < 0.01, ***P < 0.001, and no correction for multiple comparisons was applied. All other quantitative claims are assessed by goodness of fit (R^2^) and information-criterion model comparison rather than by null-hypothesis testing (see below).

#### Curve fitting, confidence intervals, and model selection

Full methodology is given above under “Parameter fitting, model selection, and uncertainty quantification” and is not repeated here. In brief, per-receptor operational models were fit by non-linear least squares, joint multi-condition AC models by Nelder-Mead, and single-parameter model comparisons by bounded scalar minimization; parameter 95% confidence intervals come from a 200-iteration cell-level bootstrap (per-receptor fits) or profile-likelihood scans (joint fits); and competing models were compared by AICc together with held-out R^2^. Fitted parameters with confidence intervals are in Supplementary Tables S1 to S3.

#### Data exclusions, randomization, and blinding

Data exclusions were defined a priori and are detailed above (the 55% forskolin-ceiling truncation for Gαs-coupled fits, the 50-cell minimum per heatmap tile, and the three AC6-knockdown plates dropped from the Fig. 5D Gαi inhibition fit). No other data were excluded; sample sizes were not predetermined by a power analysis.

## Data availability

All datasets analysed in this study are publicly available: human brain single-cell transcriptomes from the CellxGene Census (release 2025-01-30; cellxgene.cziscience.com), *Drosophila* head single-cell data from the Fly Cell Atlas, and dopamine binding affinities from the IUPHAR/BPS Guide to Pharmacology (guidetopharmacology.org). All data generated in this study: raw flow cytometry (.fcs) files, gated single-cell event arrays, processed per-figure data, CHO-K1 bulk RNA-seq, and RT-qPCR data, are deposited at CaltechDATA (DOI: 10.22002/g8586-gk090).

## Code availability

All analysis code, comprising the complete processing and modeling pipeline (Python notebooks and shared modules) used to generate the figures, is deposited at CaltechDATA (same DOI: 10.22002/g8586-gk090).

Gregrowicz, J. and Elowitz, M.B. **Adenylyl cyclases combinatorially integrate opposing dopamine receptor signals**: *data and code* (CaltechDATA, 2026. DOI: 10.22002/g8586-gk090)

## Supplementary Information

## Supplementary Figures and Tables

**Figure S1.**
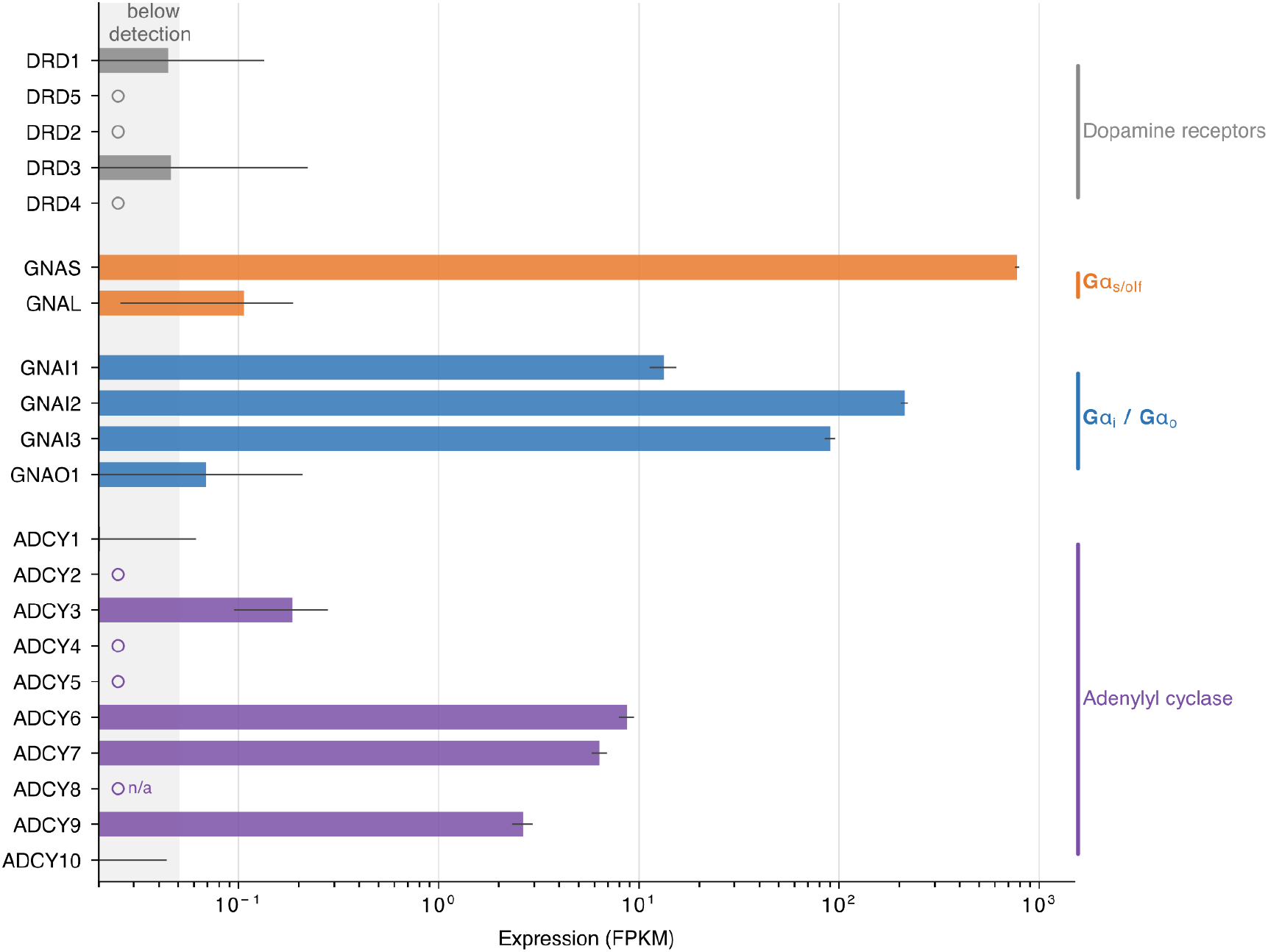
Baseline transcriptomic profile of the parental CHO-K1 cell line. Bulk RNA-seq of the wild-type CHO-K1 line characterizes the endogenous transducer and effector expression context for the heterologous signaling assay. Gene expression is quantified as FPKM (fragments per kilobase of transcript per million mapped reads) aligned to the NCBI CriGri-1.0 genome. Bars represent FPKM point estimates; error bars indicate 95% confidence intervals; open circles denote zero or sub-detection-limit values; ‘n/a’ denotes a gene absent from the reference annotation. The light grey shaded region indicates the below-detection zone (FPKM < 0.05). Full source data are provided in the CaltechDATA deposit (Data availability).

**Dopamine receptors:** All five subtypes (DRD1–DRD5) are below the detection limit, confirming CHO-K1 as a receptor-free background. **Gαs / Gαolf transducers: Gαs (GNAS) is abundantly expressed (773 FPKM), providing the stimulatory pool for DRD1- and DRD5-mediated cAMP production, whereas the neuronal Gαolf subunit (GNAL) is near the detection limit (0.11 FPKM). Gαi / Gαo transducers:** GNAS is highly expressed (773 FPKM), establishing a single dominant Gαs pool, whereas the brain-specific paralog GNAL is below detection. This large GNAS abundance supports the assumption GT ≫ KAC (Supplementary Information), yielding the Km,eff ≈ Km / [Gs] scaling validated in Fig. 3 and utilized in the operational model. The Gαi family is abundantly expressed (GNAI2 = 212, GNAI3 = 90, GNAI1 = 13 FPKM), providing the necessary transducer pool for the DRD2-mediated inhibitory responses (Figs. 3–5). Conversely, the Gαo subunit GNAO1 is near the detection limit (0.07 FPKM). Because DRD3 canonically couples to Gαo, this minimal expression accounts for the lack of a DRD3-mediated cAMP response (Fig. S3), justifying its use as a silent control (Supplementary Table S1). **Adenylyl cyclases:** The endogenous effector pool is dominated by ADCY6 (8.7 FPKM), ADCY7 (6.4), and ADCY9 (2.6). Isoforms ADCY1, ADCY2, ADCY4, ADCY5, and ADCY10 are below detection, and ADCY3 is minimally expressed (0.19 FPKM); ADCY8 is absent from the reference annotation. This RNA-seq profile of the parental line provided the initial, independent transcript-level evidence that these three isoforms dominate the endogenous pool, motivating the three-isoform integration model (Fig. 4). The isoform fractions used in that model are the reporter-line values measured by qPCR (Fig. 5A). Absolute fractions differ modestly between the two assays and cell-line states (for example, the AC7 fraction), while the rank order of the dominant AC6 is preserved.

**Figure S2.**
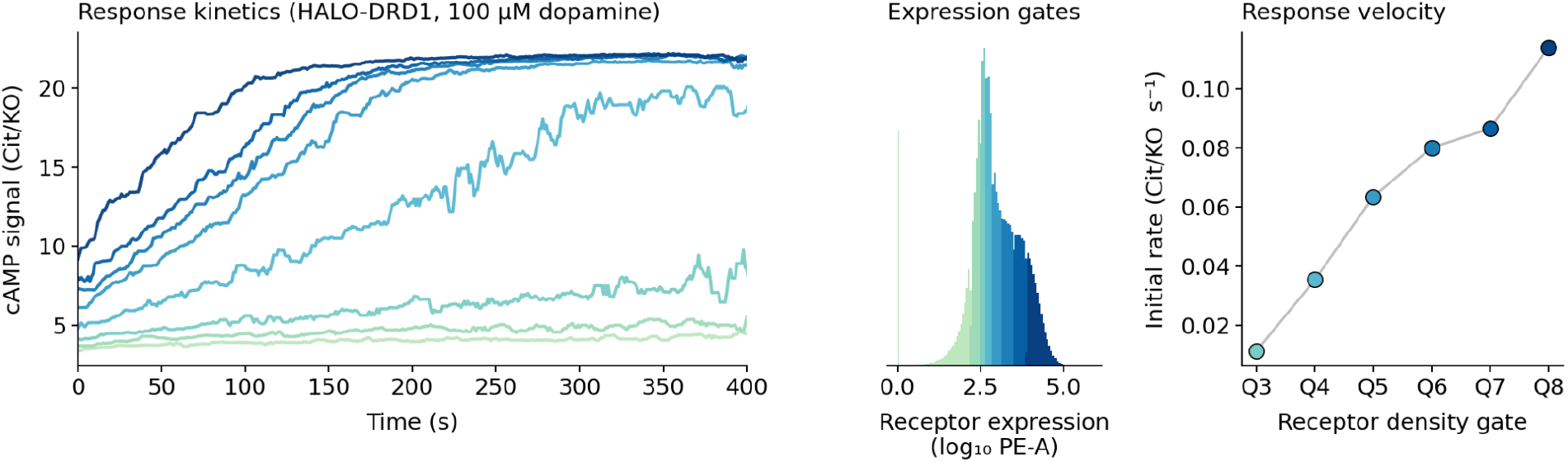
Response kinetics of cAMP signaling scale with HALO-DRD1 surface density. CHO-K1::cAMPinG1 cells transiently expressing HALO-DRD1 mRNA (Methods) were stimulated with 100 µM dopamine and continuously monitored on the CytoFlex S for ~400 s across three sequentially acquired wells (n = 1,770,189 single-cell events total). Wells were time-stacked into a single continuous timeline by zeroing each well’s clock to its first event and offsetting by the cumulative duration of the previous wells (5 s gap between wells). (A) Per-quantile median cAMP signal (Cit/KO ratio) versus time. Cells were partitioned into eight quantile gates (Q1–Q8) of HALO-DRD1 surface intensity (PE-A channel); each trace is the median of cells in that gate over 1 s time bins, smoothed with a 10 s rolling median. Quantile is encoded by the GnBu colormap, light (Q1, lowest expression) to dark (Q8, highest expression). The reporter background (Cit/KO ≈ 3.5) and the forskolin-saturated ceiling (Cit/KO ≈ 22) bracket the dynamic range. (B) Histogram of log_10_ HALO-DRD1 surface intensity, banded by quantile gate using the same colors as panel A. The eight bands span approximately three decades of receptor surface density (~10^2^ to ~10^5^ PE-A units). (C) Initial response velocity per quantile gate, computed as the slope of a linear fit on the 0–80 s window of each smoothed trace from panel A. Initial rate scales monotonically with HALO-DRD1 density and spans roughly an order of magnitude across the upper six quantile gates (Q3 = 0.011, Q4 = 0.035, Q5 = 0.063, Q6 = 0.080, Q7 = 0.086, Q8 = 0.114 Cit/KO units per second). Q1 and Q2 are omitted because their kinetic traces are indistinguishable from reporter baseline; the lowest-expressing cells produce no measurable response on this timescale.

**Figure S3.**
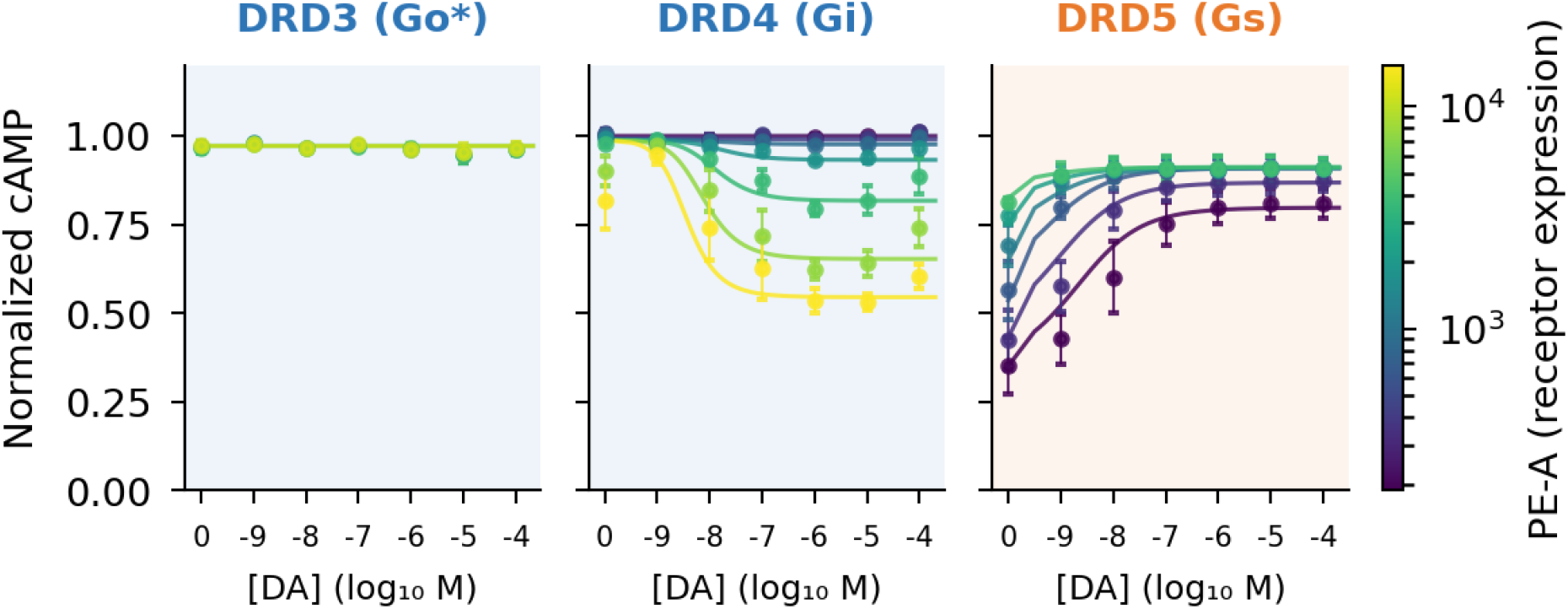
Dopamine dose-response of the remaining receptor subtypes DRD3, DRD4, and DRD5. (A–C) Single-cell dose-response curves for (A) Gαo-coupled DRD3, (B) Gαi-coupled DRD4, and (C) Gαs-coupled DRD5 in CHO-K1::cAMPinG1 cells transiently expressing HALO-tagged receptors. The Gαi/o-coupled subtypes (DRD3, DRD4) were stimulated in the presence of 10 µM forskolin to elevate the baseline cAMP level; the Gαs-coupled DRD5 was measured from the unstimulated baseline. Cells are binned and colored by their measured surface receptor abundance (arbitrary flow-cytometry units; see color bar). Solid lines represent fits of the operational model with binding affinity (Kd) locked to established IUPHAR values (fit values and 95% CIs are provided in Table S1). Data represent per-bin medians, with error bars indicating ±1 SEM across independent replicate dates (n = 2 for DRD3, n = 3 for DRD4, n = 4 for DRD5). (A) DRD3 produces no dopamine-evoked cAMP response, because its primary transducer Gαo is not endogenously expressed in CHO-K1 (Fig. S1); it therefore serves as a silent control. (B) DRD4 shows shallow, expression-dependent suppression of the forskolin-elevated baseline (R^2^ = 0.91), consistent with DRD4’s canonical preference for Gαo over Gαi. (C) DRD5 shows expression-dependent cAMP stimulation (R^2^ = 0.97). These subtypes are reported for completeness of the receptor panel and are not used in any quantitative downstream analysis; the identifiability of their fits is examined in Figure S7.

**Figure S4.**
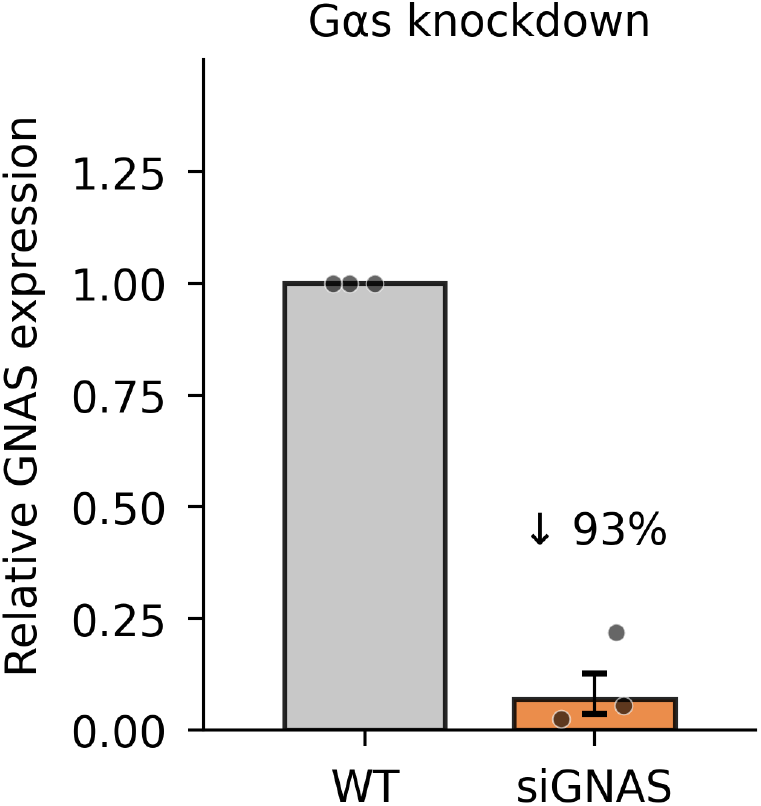
siRNA knockdown of GNAS (Gαs). RT-qPCR validation of Gαs depletion in CHO-K1::cAMPinG1 cells transfected with non-targeting siRNA (siNT, reference = 1.0) versus GNAS-targeting siRNA (siGNAS, 33 nM). siGNAS reduces GNAS mRNA by 93% (geometric-mean fold change = 0.068; per-run knockdown 78–97%). Bar, geometric mean; dots, per-run fold changes (n = 3 independent runs); error bar, ±1 log_2_ SEM. GAPDH was the reference gene; fold changes are ΔΔCt relative to the per-run siNT condition. Given the small sample (n = 3), we report the size of the knockdown rather than statistical significance; the one-sample t-test on log_2_ fold change is borderline (t = 4.29 versus the df = 2 two-tailed 5% threshold of 4.30; p = 0.05) despite the large, consistent reduction. This knockdown level is the one used to estimate the available Gαs pool in the Figure 3 G-protein-perturbation analysis.

**Figure S5.**
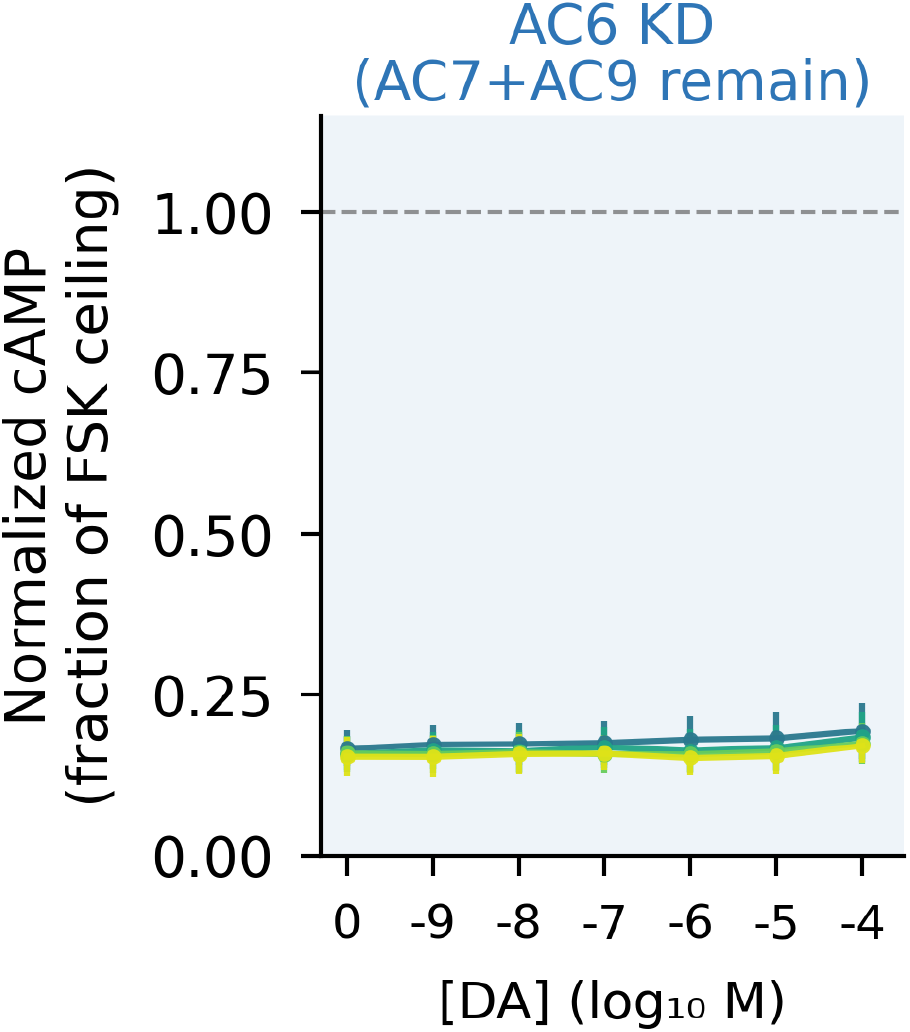
AC6 is the only forskolin-responsive adenylyl cyclase in CHO-K1 cell line. Single-cell DRD2 dopamine dose-response measured against a 10 µM forskolin baseline in AC6 knockdown (siAC6; AC7 and AC9 remaining) CHO-K1::cAMPinG1 cells, binned and colored by surface DRD2 abundance (arbitrary flow-cytometry units; see color scale). cAMP is shown on the absolute normalized scale (0 = reporter background, 1 = forskolin ceiling; dashed line). Knockdown of AC6 abolishes the forskolin response: cAMP remains flat near baseline (~0.17 of the ceiling) across all dopamine concentrations and all DRD2 expression levels, even though AC7 and AC9 are still present. By contrast, AC6-containing cells reach roughly two-thirds of the ceiling and show dopamine-dependent DRD2 inhibition (wild-type; see Figure 5D). Data are per-bin medians with error bars indicating ±1 SEM. This identifies AC6 as the sole forskolin-responsive isoform underlying the DRD2 Gαi-inhibition measurements in Figure 5D.

**Figure S6.**
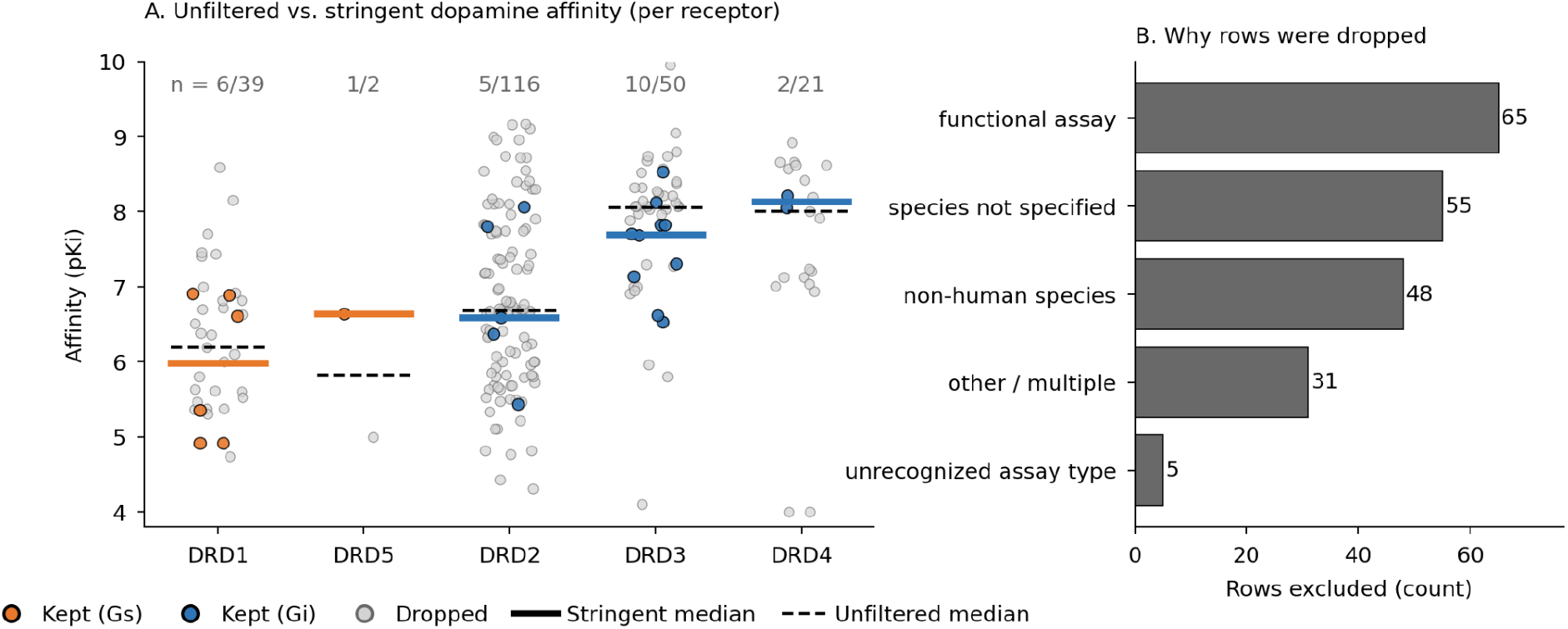
Curation of dopamine binding affinities: unfiltered database scatter and exclusion-reason breakdown. (A) Per-receptor scatter of dopamine binding measurements before and after the stringent curation that produced Figure S6. Each colored point (orange = Gαs-coupled, blue = Gαi-coupled) is one of the 24 measurements retained in the main figure; each light-grey point is a measurement excluded by one or more curation criteria (see Methods and panel B of this figure). All rows that map to one of the five dopamine receptors are included, with descriptions like “D2L”, “D2A”, and “D2long” mapped to DRD2 (long isoform) by an extended description-matching rule applied in this SI figure but not in the main-text curation, in order to expose the full per-receptor scatter that the database contains. n (kept/total): DRD1 = 6/39, DRD5 = 1/2, DRD2 = 5/116, DRD3 = 10/50, DRD4 = 2/21. Solid colored bars, stringent median (matches Figure S6); dashed black ticks, unfiltered median across all rows shown. The two medians lie within 0.1 log unit (≈25%) of each other for every receptor that has more than one entry, indicating that the stringent filter narrows the database but does not systematically bias the central tendency. (B) Counts of GtoPdb rows excluded by each curation criterion. Each row is assigned to the first applicable category in the order: functional assay (Assay Type = “F” in GtoPdb; signaling readouts such as cAMP, β-arrestin recruitment, or [^35^S]GTPγS binding rather than direct ligand binding), species not specified (assay description does not explicitly state the species), non-human species (description names rat, mouse, bovine, dog, or porcine receptor), other / multiple (combinations of the above or unparseable values, value reported as a range, or “unknown origin” phrase in description), unrecognized assay type (Assay Type column does not contain the standard B/F values).

**Figure S7.**
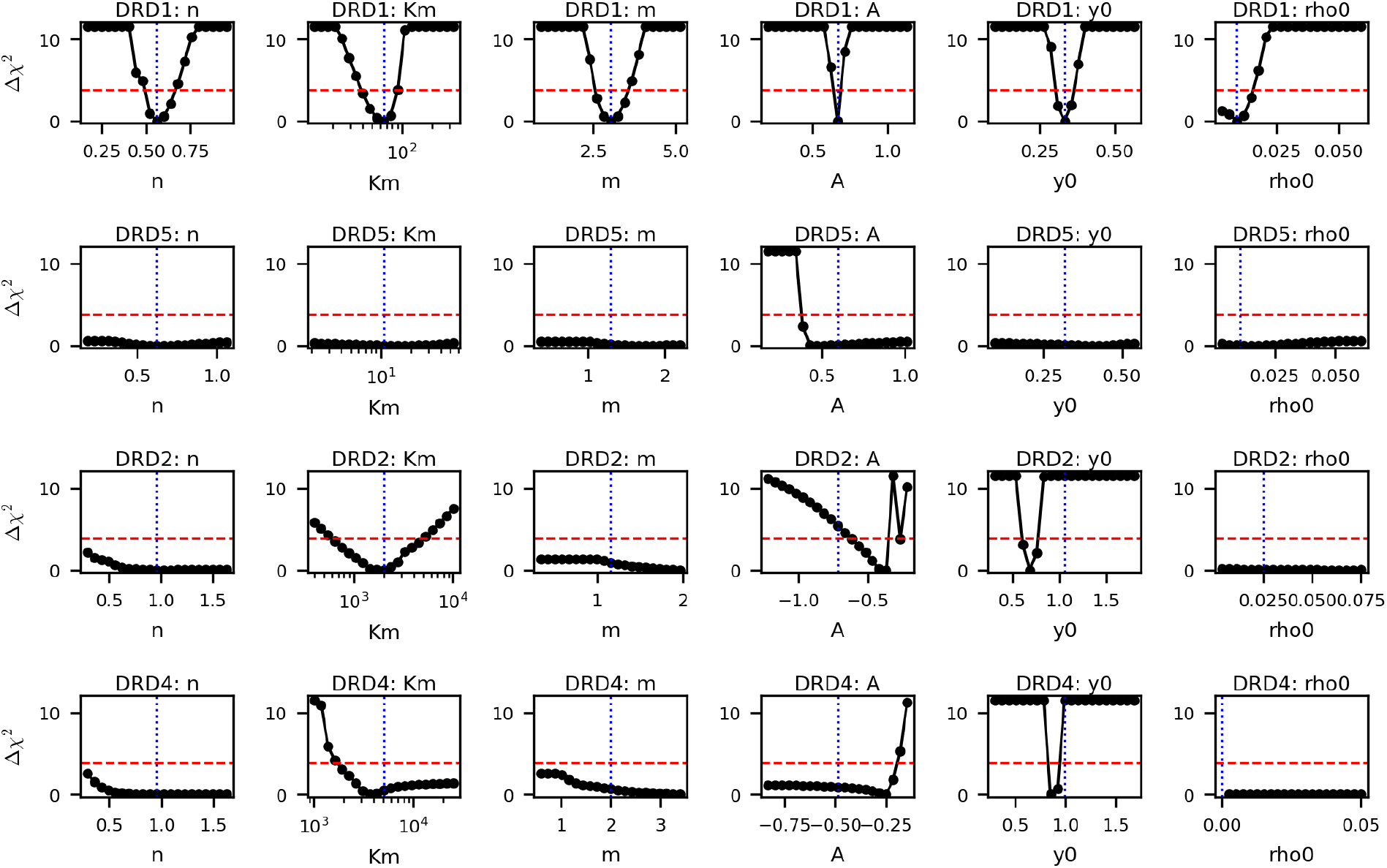
Profile-likelihood identifiability check for the per-receptor operational-model fits of Figures 3 and S3. For each well-fit dopamine receptor (rows: DRD1, DRD5, DRD2, DRD4) and each of the six fitted parameters of the modified operational model (columns: *n, K*_*m*_, *m, A, y*_0_, ρ_0_; Methods and Appendix Eq. A1.20), one parameter was held fixed on a 21-point grid spanning ±0.7 fractional units (or ±1 log unit, log-spaced, for *K*_*m*_) around the panel-A point estimate, and all remaining parameters were refit using the canonical Figures 3 and S3 pipeline. Each subpanel plots 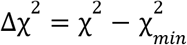 as a function of the pinned parameter. Blue dotted line, panel-A point estimate; red dashed line, the Δχ^2^ = 3. 84 critical value defining the 95% profile-likelihood confidence interval (α = 0. 05, 1 d.o.f.). Δχ^2^ is capped at 11.5 in the plot so the near-minimum shape remains visible across all panels. A sharp V-shaped profile means the parameter is well-determined; a flat or one-sided profile means the data cannot uniquely pin the parameter. The 95% profile-likelihood interval does not assume a Gaussian sampling distribution and is therefore complementary to the cell-level bootstrap intervals reported in Supplementary Table S1.

### DRD1 (top row)

All six parameters show clean V-shaped minima centered on the panel-A point estimate, confirming identifiability. Bootstrap and profile intervals agree to within a factor of two for *n, K*_*m*_, *m*, and ρ_0_. The *A* and *y*_0_ profiles are very narrow (sharp V’s), reflecting that the amplitude and baseline are tightly constrained by the data.

### DRD2 (third row)

*K*_*m*_ shows a textbook V-shaped minimum centered at ~2,000 PE-A units × occupancy: the headline parameter for the receptor-pool integration story is well-determined and the profile and bootstrap intervals coincide. *n* and *m* are bounded from below (the data require *n*≳0. 5 and *m*≳1) but flat above their point estimates, which is honest about a Hill-cooperativity claim we are not making; the qualitative conclusion that DRD2 occupancy and pool transduction are approximately non-cooperative remains supported. The amplitude *A* and baseline *y*_0_ profiles place the chi-square minimum at slightly different values than the canonical fit, which optimizes a robust soft-L1 loss rather than the raw χ^2^. The profile shape and width remain a valid identifiability readout in either case, and the offset reflects an internal correlation between the Gαi-suppression amplitude and the forskolin-stimulated reference baseline that is unique to Gαi-coupled receptors (the readout is anchored to a separately stimulated ceiling rather than a calibrated zero) and that does not affect the Figure 3 perturbation analysis or the receptor-comparison narrative. The constitutive activity ρ_0_ profile is flat: DRD2 constitutive activity is not separable from forskolin-ceiling drift in this assay, and the parameter was fit within physical bounds [0, 0.5].

### DRD5 (second row)

All six profiles are flat. With four replicate dates and only two valid expression bins after the PE-A threshold cutoff, the dataset is underdetermined for the six-parameter operational model: multiple parameter combinations fit the visible data equally well, and the optimizer always finds a near-baseline χ^2^ regardless of where in the grid one parameter is pinned. The point estimates remain consistent with the data, they are simply not uniquely identifiable. Importantly, the *shape* of the DRD5 dose-response curve in Figure S3 is qualitatively as expected for a Gαs-coupled receptor (positive amplitude, monotonic rise, saturating plateau), and the assay successfully detects the receptor-driven signal above background. DRD5 is reported in Figure S3 for completeness of the receptor panel (DRD1–DRD5) and as a positive control demonstrating that the platform reports Gαs signaling from a second dopamine-receptor subtype; it is **not used in any downstream analysis**. Figures 3–5 use DRD1 with and without Gαs perturbations, DRD2 single-receptor Gαi titrations, and DRD1 + DRD2 combinatorial pairings only.

### DRD4 (bottom row)

Similar identifiability pattern to DRD5: *K*_*m*_ shows a wide U-shaped minimum near the canonical ~5,000 PE-A units × occupancy (point inside the 95% interval), but the CI spans roughly an order of magnitude; *n, m, A*, and ρ_0_ profiles are flat or bounded only from one side. This is the expected behavior for a receptor whose canonical primary transducer (Gαo) is absent from CHO-K1: DRD4 in this background couples through a residual Gαi pool with shallow dynamic range, which limits the data’s capacity to disambiguate Hill slope from pool cooperativity. The dose-response shape in Figure S3 is again qualitatively consistent with a Gαi-coupled signal (negative amplitude, monotonic suppression below the forskolin ceiling), confirming that the assay reports DRD4-driven signaling. As with DRD5, DRD4 is **not used in any downstream analysis** and is reported here as a positive control documenting the assay’s ability to detect a fourth dopamine-receptor subtype on a non-canonical transducer.

### Summary

The headline identifiability conclusion supports the use of DRD1 and DRD2 fit parameters in downstream Figures 3–5: *K*_*m*_ is well-determined for both receptors, and the bootstrap and profile-likelihood intervals agree where it matters. DRD3 (Gαo-coupled, silent in CHO-K1; not shown), DRD4, and DRD5 are reported in Figure S3 to document the full pharmacology of the platform but are not relied on for any quantitative downstream claim. Source code, per-profile CSVs, and the figure are reproducible from the deposit; profiles use the same data extraction, gating, and contamination correction as the canonical Figures 3 and S3 fit (Methods).

**Figure S8.**
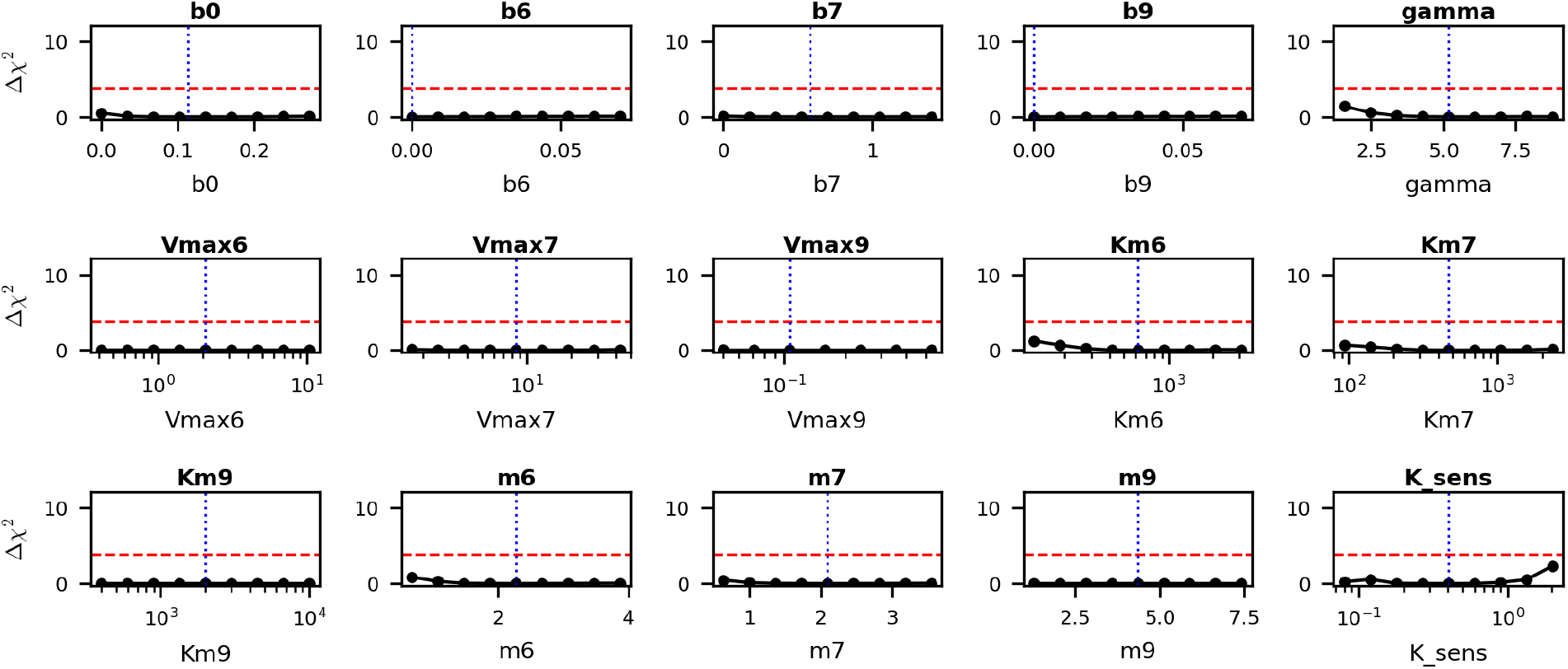
Profile-likelihood identifiability check for the Figure 5 joint per-AC fit. For each of the 15 fitted parameters of the joint per-AC integration model (b_0_, b_6_, b_7_, b_9_ basal terms; *V*_*max,i*_ / *K*_*m,i*_ / *m*_*i*_ Hill parameters per isoform AC6/AC7/AC9; γ activity retention exponent; *K*_*sens*_ biosensor saturation constant), one parameter is held fixed on a 9-point grid spanning ±0.7 fractional units (linear scale) or ±1 log unit (log scale) around its point estimate, and the remaining 14 parameters are refit using the same cost function as the Figure 5 joint per-AC fit. Each subpanel plots Δχ^2^ = χ^2^ − χ^2^_min as a function of the pinned parameter; blue dotted line, point estimate; red dashed line, Δχ^2^ = 3.84 critical value (95% confidence cutoff at α = 0.05, 1 d.o.f.).

All 15 profiles remain below the cutoff across the full grid range. Wider sweeps on Vmax_6_, *K*_*m*,6_, and γ (span 1.5 = 3 log decades; not shown) similarly fail to bracket finite confidence intervals. The cost surface is flat across 1–2 orders of magnitude in most parameter directions, indicating that the joint fit is practically identifiable (the optimization converges to a finite χ^2^_min) but individually non-identifiable at the per-parameter level: many distinct parameter sets fit the data equally well^40^.

The model’s predictive power, demonstrated by the parameter-free dual-AC predictions of Figure 5E (R^2^ = 0.94–0.97 on the held-out AC6+AC7, AC6+AC9, and AC7+AC9 contexts), reflects identifiable parameter combinations rather than identifiable individual parameters. We interpret the model accordingly as evidence that the AC isoform layer with its assumed Gs/Gi/Gβγ wiring is sufficient to reproduce the observed combinatorial integration logic, not as a quantitative inventory of per-isoform biochemical constants.

**Figure S9.**
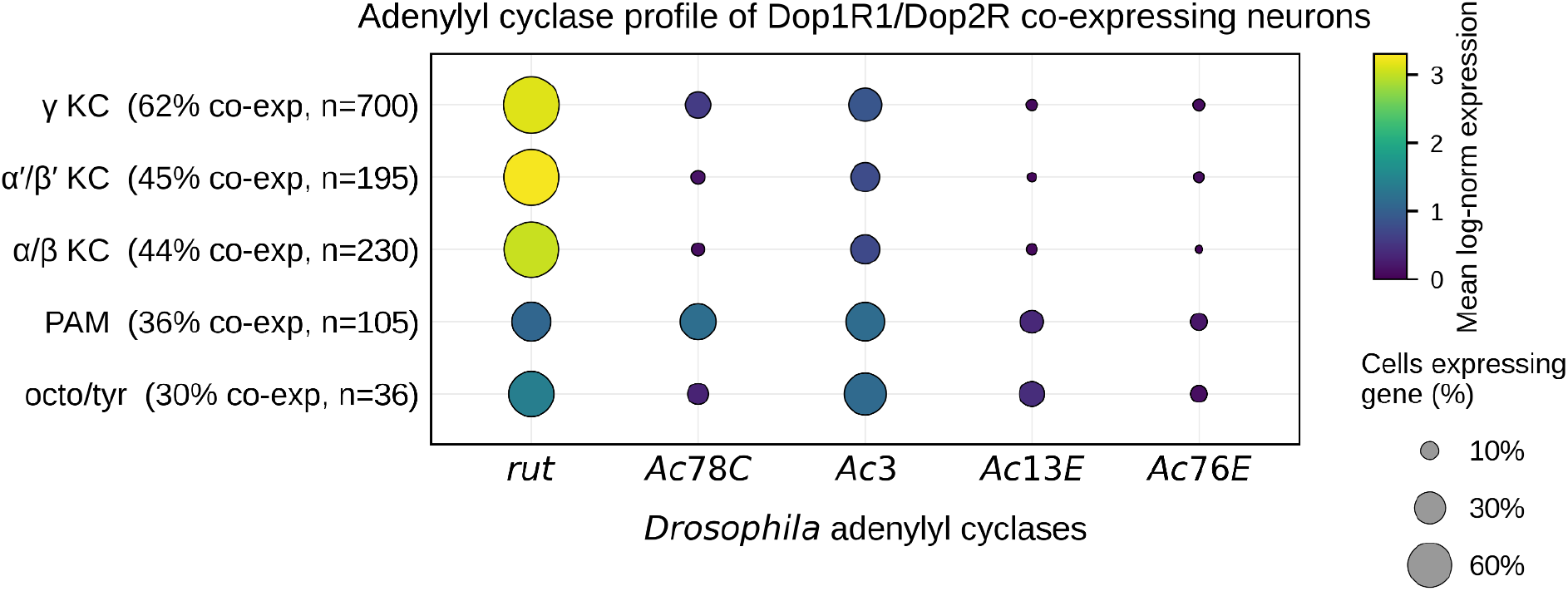
Adenylyl cyclase expression in Dop1R1/Dop2R co-expressing *Drosophila* neurons. Dot plot of the five adenylyl cyclases (rut, Ac78C, Ac3, Ac13E, Ac76E) in cells co-expressing the D1-like receptor Dop1R1 and the D2-like receptor Dop2R (both with raw count > 0), across *Drosophila* head cell types (Fly Cell Atlas, head 10x): three Kenyon-cell subtypes (γ, α′/β′, α/β), dopaminergic PAM neurons, and octopaminergic/tyraminergic neurons. Dot size, percentage of co-expressing cells with detectable transcript (raw count > 0); colour, mean log-normalized expression (counts normalized to 10,000 per cell, log1p). Row labels give each cell type’s co-expression percentage and the number of co-expressing cells.

**Figure S10.**
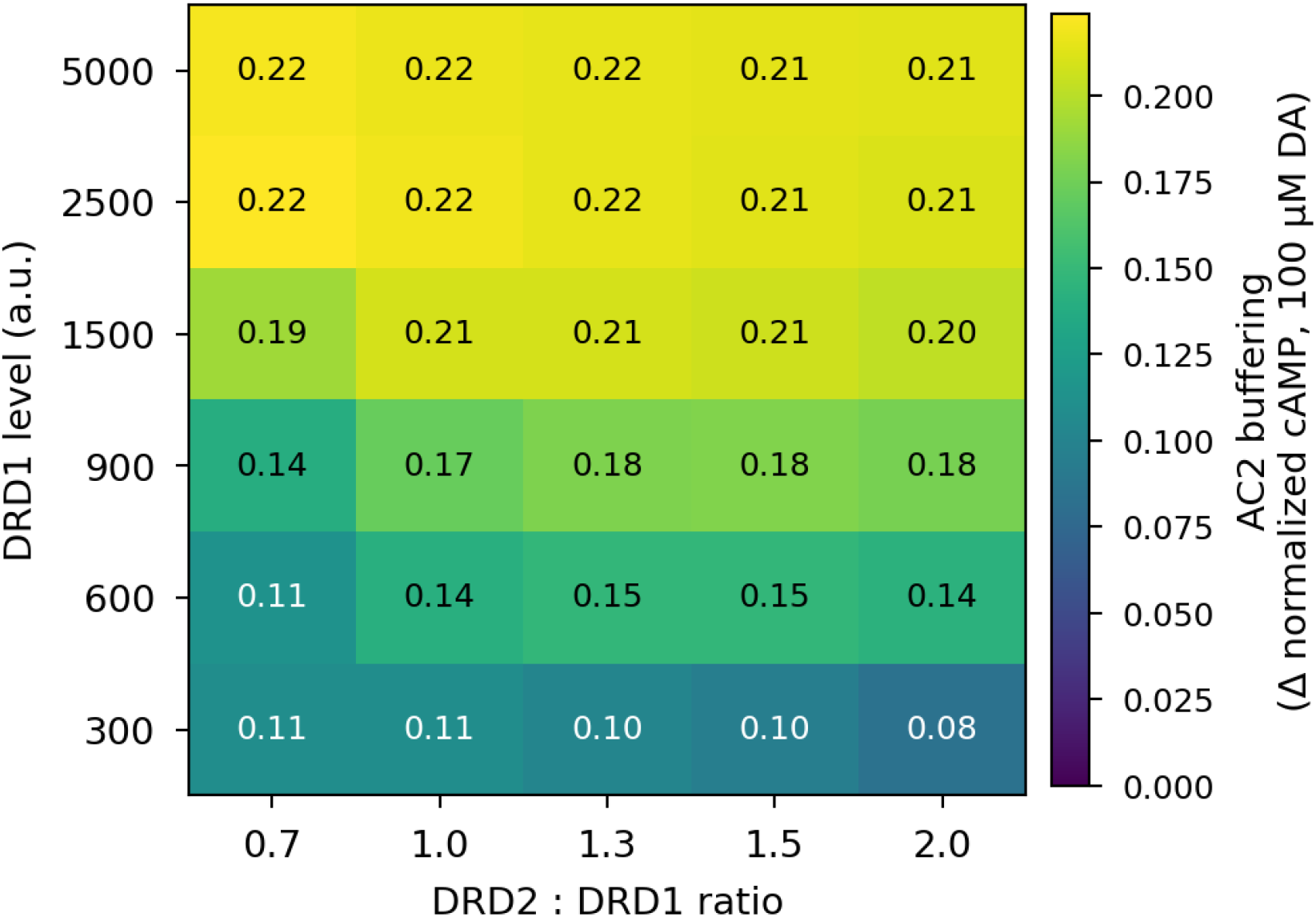
Simulated eMSN AC2 buffering is robust to receptor level and DRD2:DRD1 ratio. All values are simulations of the fitted operational AC-integration model (as in Fig. 6D), not measurements. AC2 buffering is defined as the difference in simulated normalized cAMP between the wild-type eMSN (AC2 at the eMSN level) and the same simulated cell with AC2 reduced to the canonical dMSN level, computed at saturating dopamine (100 µM) across a grid of absolute DRD1 surface levels (300–5000 a.u.) and DRD2:DRD1 ratios (0.7–2.0). Normalized cAMP is expressed relative to the forskolin-saturated maximum (0 = reporter baseline, 1 = saturation). In the model, buffering is positive in every condition (0.08–0.22) and increases with receptor level and Gαi tone, so the wild-type eMSN sustains more cAMP than the AC2-reduced profile across the simulated range of receptor levels (in absolute terms the wild-type response spans 0.25–0.63 and the AC2-reduced response 0.13–0.52). The receptor levels used in Fig. 6D (≈1500 a.u.; ratios 1.0–1.3) lie in the middle of this grid, indicating that the simulated eMSN result is not an artifact of the chosen receptor levels.

**Figure S11.**
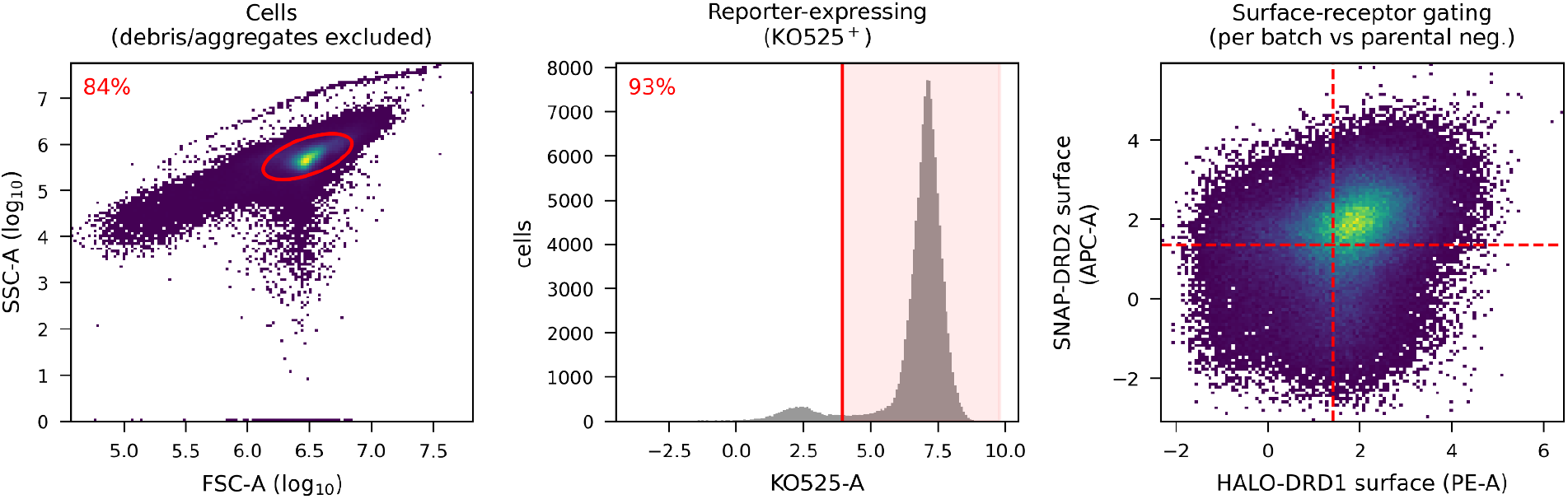
Representative flow cytometry gating strategy. Sequential gating of a representative combinatorial (HALO-DRD1 + SNAP-DRD2) sample. Left: FSC-A × SSC-A gate excluding debris and aggregates. Middle: reporter-expressing gate on KO525 (defined as the central 99% of the KO525 distribution of parental CHO-K1::cAMPinG1 cells; Methods). Right: surface-receptor gating on the HALO-DRD1 (PE-A) and SNAP-DRD2 (APC-A) surface-label channels, with thresholds set once per cell-line batch against the parental label-negative line. Percentages are the fraction of events retained at each step for this example. FSC-A/SSC-A, log_10_ scale; fluorescence, arcsinh scale.

**Supplementary Table S1.** Per-receptor operational-model fit parameters (Figures 3 and S3). Parameters of the modified operational model (Appendix Eq. A1.20) fitted to the binned single-cell dose-response data of Figures 3 and S3. Values are point estimates with 95% confidence intervals from a 200-iteration cell-level empirical bootstrap (cells resampled with replacement within each (date × dopamine concentration) sample, per-bin medians recomputed by re-binning the resampled cells on the surface-receptor channel, and the model refit through the same Trust Region Reflective + soft-L1 pipeline); *K*_*d*_ values are locked at the IUPHAR/BPS Guide to Pharmacology median per receptor (Fig. S6) and not fitted. Complementary profile-likelihood scans on the same parameters are shown in Supplementary Figure S7 and tabulated in the deposit as an identifiability cross-check; the bootstrap and profile intervals agree to within a factor of two for the well-fit DRD1 and DRD2 receptors. The Dates column counts independent experimental dates per receptor and is therefore a lower bound on the number of biologically independent replicates: multiple wells acquired on a single date also represent biologically independent replicates and are not separately enumerated here.

| Receptor | Coupling | Kd locked (nM) | Km (nM) | *n* | *m* | *A* | R<sup>2</sup> | Dates |
| --- | --- | --- | --- | --- | --- | --- | --- | --- |
| DRD1 | Gs | 2,356 | 65 [20, 122] | 0.56 [0.42, 0.85] | 3.0 [1.8, 4.3] | 0.67 [0.60, 0.76] | 0.98 | 4 |
| DRD5 | Gs | 229 | 11 [2, 38] | 0.62 [0.43, 0.92] | 1.3 [1.0, 2.3] | 0.59 [0.44, 0.75] | 0.97 | 4 |
| DRD2 | Gi | 257 | 2,000 [1,510, 2,690] | 0.96 [0.70, 1.31] | 1.2 [1.0, 1.7] | -0.72 [-0.82, -0.62] | 0.97 | 5 |
| DRD4* | Gi | 8 | ≈ 5,000 | ≈ 1.0 | ≈ 2.0 | ≈ -0.49 | 0.92 | 3 |
| DRD3 | Gao | 20 | n.f. | n.f. | n.f. | n.f. | n.f. | 2 |

\*DRD4: bootstrap CIs for Km, *n*, *m*, and *A* all reach the optimizer bounds (CIs span more than five log units), reflecting the limited number of replicate dates ( $n = 3$ ) and the modest dynamic range of DRD4's Gai-coupled response in CHO-K1, consistent with DRD4's canonical preference for Gao (absent in CHO-K1) over Gai. Point estimates are reported without CIs; the fit converges but should not be interpreted as a tight pharmacological measurement. n.f. = not fitted. DRD3 signals through Gao, which is absent from CHO-K1; the dose-response is flat at reporter baseline and the operational model is not estimable.

### Shared receptor parameters (all conditions)

**Supplementary Table S2.** Shared-parameter fit for DRD1 ± Gαs perturbation (Figure 3C,D). The receptor-intrinsic parameters (*n, K*_*m*_, *m, A, y*_0_, ρ_0_) are shared across all conditions; only the transducer pool [*Gs*] differs per condition. Wildtype anchors [*Gs*] = 1. 0; the two ΔGs conditions fit a single multiplicative rescaling. Values are point estimates with 95% CI from cell-level bootstrap. AIC favors this Km-only-scaling fit (receptor-intrinsic parameters shared, [*Gs*] varying) over a both-scale alternative (in which the saturating amplitude *A* is also rescaled per condition) by Δ*AIC* = 30. 1.

Shared receptor parameters (all conditions).
| Parameter | Value [95% CI] |
| --- | --- |
| Kd (nM) | 2,356 (locked from IUPHAR median, Fig. S6) |
| *n* | 0.6 [0.4, 0.9] |
| $K_D$ (PE-A units × occupancy, in dopamine concentration units, nM) | 70 [40, 110] |
| m | 2.8 [1.7, 4.2] |
| A | 0.71 [0.65, 0.78] |
| $y_0$ | 0.21 [0.16, 0.27] |
| $\rho_0$ | 0.01 [0.0, 0.03] |
| $R^2$ (combined fit) | 0.98 |

**Supplementary Table S3.**
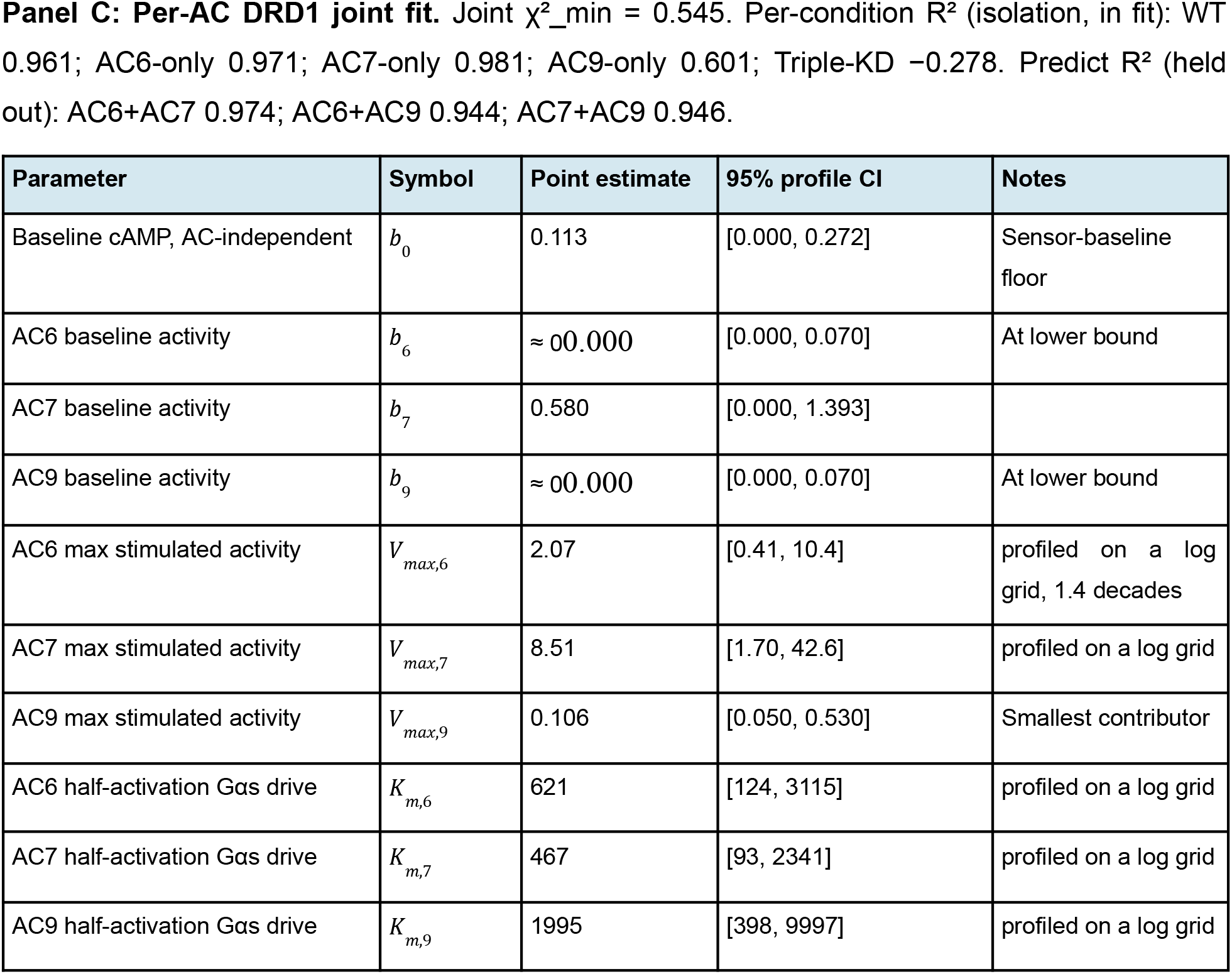
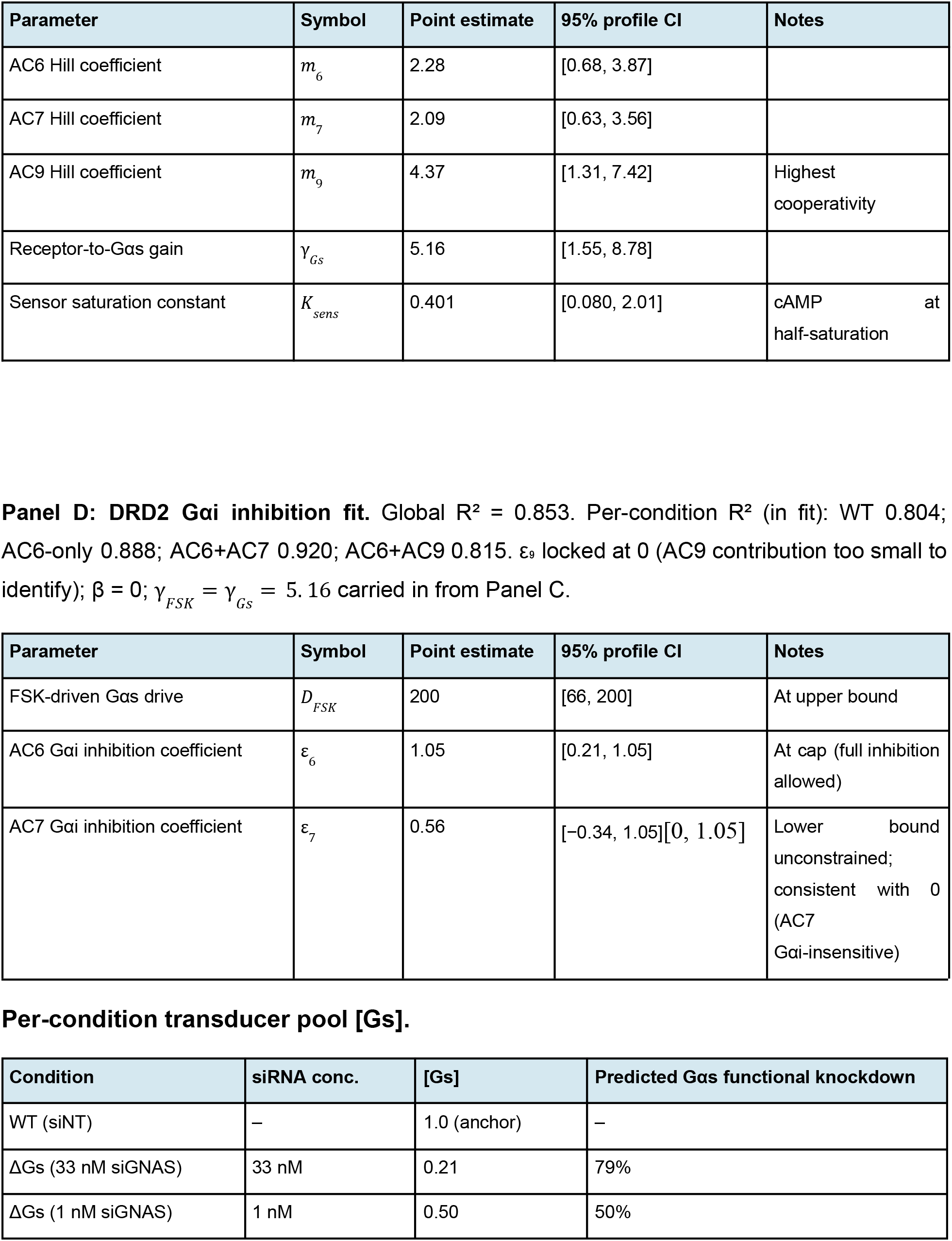
Joint per-AC fit and DRD2 Gαi inhibition fit parameters (Figure 5C,D). Free parameters of the joint per-AC operational fit (Panel C; 15 parameters fitted simultaneously across the five isolation conditions: WT, AC6 only, AC7 only, AC9 only, Triple-KD) and of the DRD2 Gαi-inhibition fit (Panel D; 3 parameters with ε_9_ = 0 locked on identifiability grounds). Profile 95% confidence intervals from one-parameter-at-a-time pinned refits with Δχ^2^ = 3.84 cutoff^41^. The breadth of per-parameter intervals reflects a practically-identifiable-but-individually-non-identifiable regime: the joint cost surface has a finite minimum (the model fits the data) but is shallow along most single-parameter directions. Predictive validity for Panel C is established independently by the held-out dual-knockdown contexts of Figure 5E (R^2^ = 0.94 to 0.97). Locked parameters (Panel C): *K*_*d*_, n, and ρ_0_ for DRD1 carried in from Supplementary Table S1; AC isoform fractional weights *F*_*i*_ (cond) derived from siRNA-validated knockdown depths and FPKM (Methods). Locked parameters (Panel D): ε_9_ = 0, β = 0, γ_*FSK*_ = γ_*Gs*_ from Panel C.

**Supplementary Table S4.**
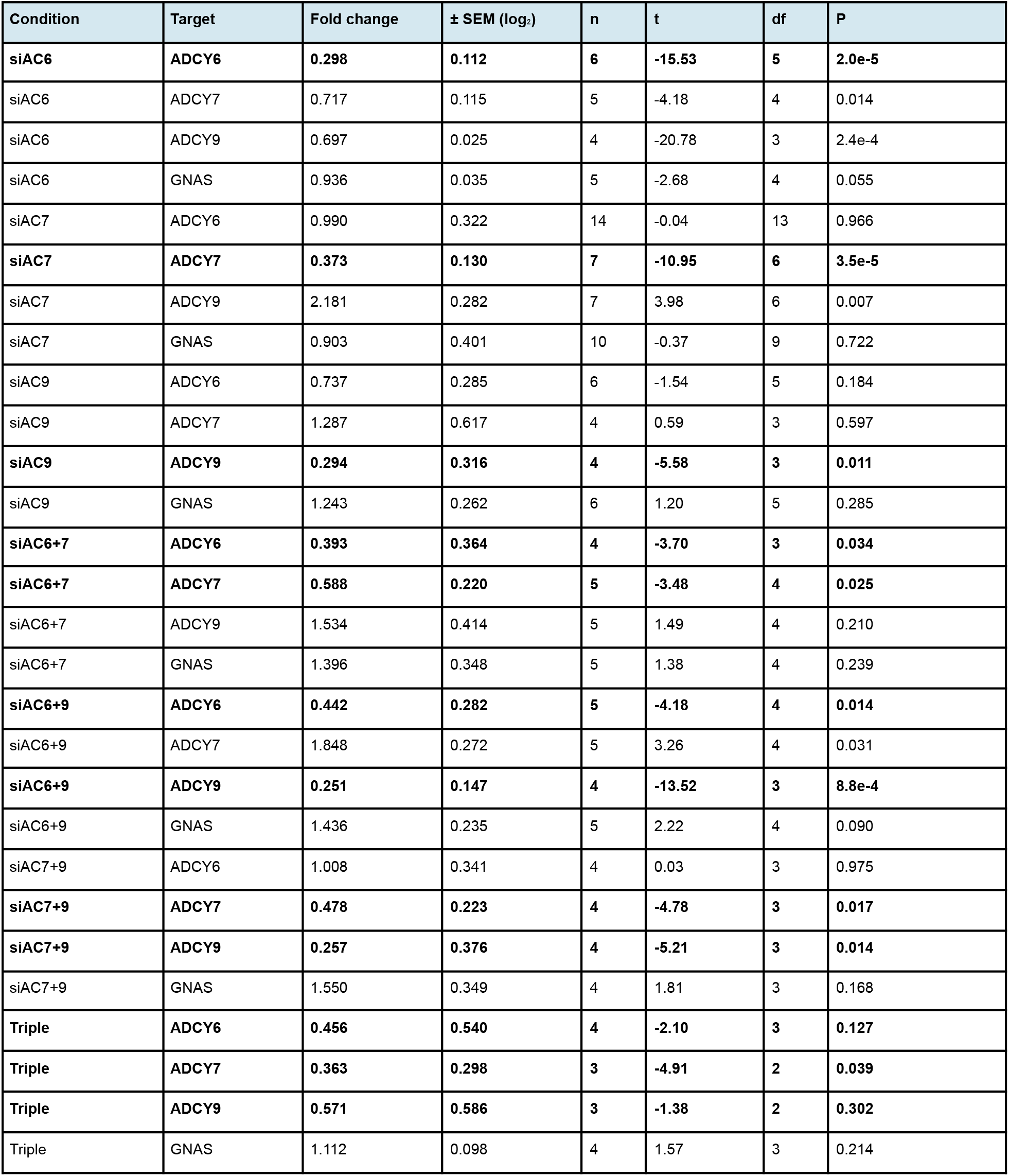
RT-qPCR knockdown efficiency and specificity (Fig. 5B). Geometric-mean fold change of each adenylyl cyclase transcript (ADCY6, ADCY7, ADCY9) and of the Gαs transcript GNAS relative to the non-targeting control (reference = 1.0), across single, double, and triple siRNA conditions. P values are from a one-sample, two-tailed t-test of the per-run log_2_ fold change against 0 (degrees of freedom = n − 1); error is ±1 log_2_ SEM. This matches the test convention used for the Gαs knockdown in Fig. S4. Bold rows indicate the transcript intentionally targeted in each condition.

### Key Resources and Tables

**KEY RESOURCES TABLE**

Key Resources and Tables KEY RESOURCES TABLE
| REAGENT or RESOURCE | SOURCE | IDENTIFIER |
| --- | --- | --- |
| **Software and algorithms** |  |  |
| Python 3.13.5 | python.org | N/A |
| Affinity Designer | Serif | v2 |
| EasyFlowQ | github.com/ym3141/EasyFlowQ/releases/tag/v1.5.6 |  |
| **Deposited data** |  |  |
| CellxGene Census | Chan Zuckerberg Initiative | v2025-01-30 |
| GproteinDb consensus coupling data | gpcrdb.org | Accessed 2025-01 |
| GtoPdb binding affinity data | IUPHAR/BPS Guide to Pharmacology | Accessed 2026-01-13 |

## Appendix 1: Derivation of the Extended Operational Model

Simple one-to-one coupling between ligand binding and cellular output is rare. In some cases, ligand-gated ion channels are one of the exceptions: the channel is itself the effector, and the ion current is a linear readout of channel occupancy,

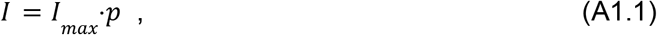

where *p* is the fractional occupancy of the channel by the ligand,

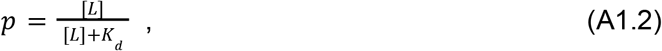

where *K*_*d*_ is the dissociation constant.

In G-protein-coupled receptor (GPCR) cascades, ligand binding sits several layers upstream of the effector. It initiates a chain of saturable steps, each one serving as the enzymatic substrate for the next, and the composition of these steps decouples ligand affinity from downstream output. The operational model resolves this decoupling phenomenologically, capturing the input-output relationship of the cascade as a composition of saturable functions.

### Layer 1: Receptor occupancy and constitutive activity

The first stage of the signaling cascade is the transition of the receptor to the active conformation (*R*^*^). Because ligand binding and receptor conformational changes reach equilibrium on a timescale of seconds, well before the ~300 second timescale required for cAMP accumulation to equilibrate in the presence of IBMX, we apply the quasi-steady-state approximation to the binding step. Rather than solving a full four-state thermodynamic isomerization cycle, the model employs a phenomenological mixture that enforces the physical boundary conditions of the system. Activity is driven by two parallel populations: receptors that spontaneously adopt the active state in the absence of ligand (ρ_0_) and receptors that transition to the active state upon ligand binding. Treating the ligand-bound complex (*RL*) as functionally equivalent to the active state, the total active fraction ρ([*L*]) is defined as the sum of constitutive activity and ligand-induced occupancy:

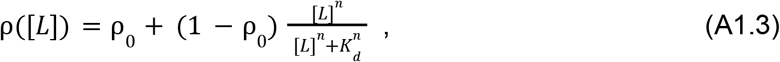

where *K*_*d*_ is the macroscopic dissociation constant and *n* is a phenomenological Hill coefficient capturing apparent cooperativity at the binding step. This expression anchors the active fraction at the constitutive floor ρ_0_ when [*L*] = 0 and approaches unity at saturating ligand concentrations. The derivation below assumes agonist ligands; for inverse agonists, an extension that permits suppression below ρ_0_ would be required.

### Layer 2: Signaling drive

Active receptors (*R*^*^) function analogously to enzymes, acting as catalysts for the signaling cascade rather than serving as the final output. Specifically, the *R*^*^ species functions as a guanine nucleotide exchange factor (GEF) on the heterotrimeric G-protein pool. The rate of G-protein activation is determined by the density of these active species at the plasma membrane. This rate-setting quantity is defined as the signaling drive *D*:

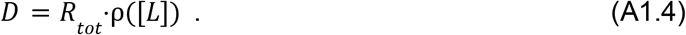

Biologically, *D* represents the concentration of active receptors available to initiate the transduction process. The per-receptor catalytic efficiency is absorbed into the downstream half-maximum *K*_*E*_. In this framework, *D* is strictly a receptor-defined quantity, whereas the availability of the transducer pool (*G*_*T*_) is explicitly modeled in the subsequent collision-coupling derivation.

### Layer 3: Collision coupling and G-protein saturation

The mapping from signaling drive *D* to the fraction *S* of activated adenylyl cyclase (AC) follows from a collision-coupling treatment of heterotrimer kinetics. Assuming that intracellular GTP concentration is not rate-limiting for exchange, the rate of change of the active G-protein pool ([*G*_*GTP*_]) is governed by the ordinary differential equation balancing activation and deactivation:

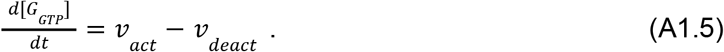

When the receptor is not saturated with respect to free G-protein (i.e., nucleotide exchange and dissociation are sufficiently fast that the receptor is not bottlenecked by substrate), the activation rate is first-order in both drive and free G-protein, and deactivation proceeds at a rate set by intrinsic GTPase activity:

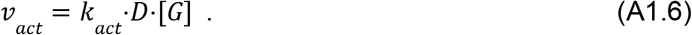

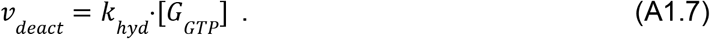

Because the G-protein GTPase cycle also equilibrates on a timescale of seconds, we apply the QSSA (*d*[*G*_*GTP*_]/*dt* = 0). Combining this condition with the conservation of mass (*G* = [*G*] + [*G*_*GTP*_]) yields a hyperbolic response of the active-transducer pool:

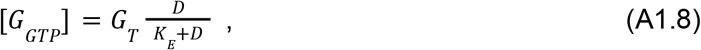

with *K*_*E*_ = *k*_*hyd*_ /*k*_*act*_. The half-maximum *K*_*E*_ depends on rate constants alone and does not scale with *G*_*T*_. Active G-protein then engages AC through saturable binding,

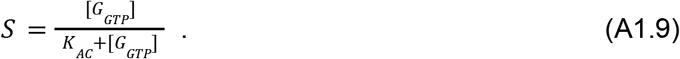

Substituting the steady-state [*G*_*GTP*_] into the AC activation function collapses the dynamic two-step chain into a single analytic transducer output,

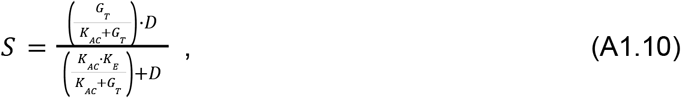

which has the form of the classical Black-Leff operational limit *S* = *S*_*max*_ ⋅*D*/(*K*_*eff*_ + *D*), with the saturating amplitude and the half-maximum of the transducer layer given by:

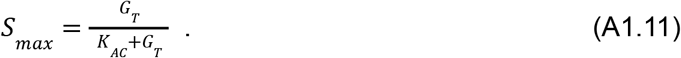

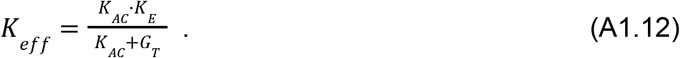

#### The *K*_*m*_ /[*G*] knockdown scaling and the GTP shift

A G-protein knockdown (e.g., via siGNAS) reduces the available pool *G*_*T*_ to a fraction α⋅*G*_*T*_. In CHO-K1 and most other mammalian systems, transcriptomic counts place G-protein abundance in vast stoichiometric excess of AC, ensuring the system operates in the *G*_*T*_ ≫ *K*_*AC*_ regime. In that limit the operational parameters collapse to:

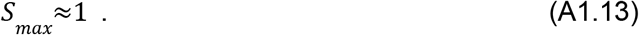

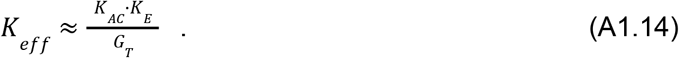

Reducing the transducer pool to α⋅*G*_*T*_ shifts the half-maximum inversely:

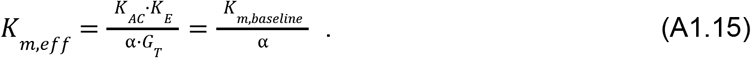

Because *S*_*max*_ = *G*_*T*_ /(*K*_*AC*_ + *G*_*T*_), as long as the depleted pool α⋅*G*_*T*_ remains much larger than *K*_*AC*_, *S*_*max*_ stays approximately 1. Consequently, the maximum achievable signal amplitude *A* remains unmoved by the knockdown, as expected analytically in the mass-action limit.

The collision-coupling limit further dictates that reducing the G-protein pool acts exclusively as a downstream substrate limitation (shifting apparent *K*_*m*_) without altering the macroscopic ligand dissociation constant (*K*_*d*_). In a live-cell assay with physiological millimolar GTP, the high-affinity ternary complex (*L*-*R*-*G*) rapidly binds GTP and dissociates. Time-averaged ligand binding is therefore dominated by the uncoupled, low-affinity receptor state, meaning the measured *K*_*d*_ is already the GTP-shifted constant. Thus, the single-parameter *K*_*m*_ /α scaling follows from the mass-action kinetics of transient receptor-G-protein coupling.

### Layer 4: System cooperativity and ultrasensitivity

The baseline collision-coupling derivation yields a standard hyperbola with a Hill coefficient of *m* = 1. However, because GPCR signaling cascades frequently exhibit steep, cooperative responses, the operational transducer function is generalized as a Hill-type sigmoid:

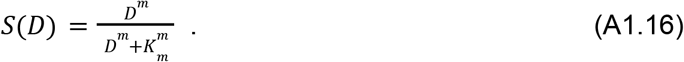

Two theoretical mechanisms can generate this active source of ultrasensitivity. The first is zero-order ultrasensitivity of the covalent G-protein activation cycle (Goldbeter and Koshland, 1981): when the opposing GEF and GAP activities both approach saturation with respect to their substrate, the steady-state fraction of active G-protein becomes switch-like, and small changes in relative velocities generate Hill coefficients well above unity. Because this regime is entered at high substrate load, zero-order ultrasensitivity is expected to strengthen with receptor density, allowing the same receptor to appear graded at low expression and switch-like at high expression.

The second mechanism is spatial compartmentalization: receptors and G-proteins confined in nanodomains or lipid rafts produce locally high concentrations where a small number of active receptors saturate their local effector pool, yielding macroscopic cooperativity in the ensemble-averaged signal. Note that the binding-step Hill coefficient *n* in (A1.3) and the transducer-step *m* here are distinct phenomenological parameters: *n* captures apparent cooperativity at ligand-receptor binding, while *m* captures downstream saturation in the G-protein cycle and effector engagement.

### Layer 5: Output mapping and biosensor saturation

The activated-transducer fraction *S*(*D*) drives the accumulation of intracellular cAMP. In the *G*_*T*_ ≫ *K*_*AC*_ regime, the maximum concentration of active G-protein is more than sufficient to fully saturate the available AC. Consequently, the maximum rate of cAMP production is bounded by the effector enzyme itself:

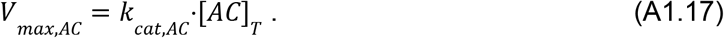

The mapping from this accumulated cAMP to the observable biosensor signal *y* introduces an additional non-linear boundary. Fluorescent cAMP biosensors are typically saturable two-state binders, where the occupancy of the cAMP binding site is coupled to a conformational shift and a resulting fluorescence change. The complete input-output relationship of the biosensor is therefore a Hill-type function:

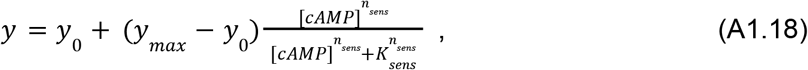

where *K*_*sens*_ is the half-saturation concentration of the biosensor, *n*_*sens*_ its phenomenological Hill coefficient, and *y*_*max*_ the absolute physical ceiling of the fluorescence ratio.

The single-pathway linear approximation. For single-receptor dose-responses and single-pathway perturbations (such as the GNAS knockdown), this full saturable mapping is unnecessary. The operational model simplifies the output to a linear approximation:

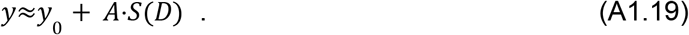

This simplification is rigorously defensible for unified signaling pathways. A cascade of sequential saturating steps (transducer activation → AC activation → biosensor engagement) can be phenomenologically collapsed into a single macroscopic Hill function. In this regime, the biosensor’s specific half-maximum (*K*_*sens*_) and cooperativity (*n*_*sens*_) do not need to be explicitly modeled; they are mathematically absorbed into the apparent transducer parameters (*K*_*m*_ and *m*) fitted in Layer 4, while the pathway amplitude *A* absorbs the linear range of the sensor.

#### The requirement for explicit sensor modeling in combinatorial architectures

While the convolution of sensor parameters into apparent *K*_*m*_ and *m* is robust for a single, unified AC pool, this mathematical shortcut breaks down when modeling combinatorial integration across heterogeneous AC isoforms.

In the multi-isoform models (Layer 6), the independent cAMP contributions from AC6, AC7, and AC9 are summed before they engage the biosensor. If one simply summed the linear approximations (*A*⋅*S*_6_ + *A*⋅*S*_7_ + *A*⋅*S*_9_), a highly co-expressing cell would yield a predicted fluorescence signal that unphysically exceeds the absolute ceiling of the biosensor (*y*_*max*_). Therefore, to accurately isolate signal integration mechanisms at the AC layer without convoluting them with downstream biosensor overflow, the combinatorial multi-isoform models must explicitly re-introduce the saturable biosensor envelope. By fixing the sensor parameters (*y*_*max*_ and *n*_*sens*_) via a direct, receptor-independent forskolin calibration, the model sets a fixed upper bound on the response. This constrains the fit to saturating, pathway-limited behavior rather than unbounded linear growth.

#### The complete composition

Substituting each layer into the next collapses the cascade into a single composite expression:

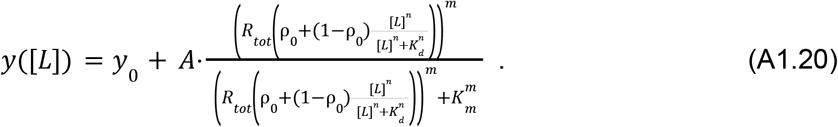

The signature of this composition is a pair of nested saturating functions, one for occupancy and one for transducer activation, separated by the expression-dependent drive *R*_*tot*_ ⋅ρ([*L*]). Theexistence of these two distinct biochemical thresholds allows a single receptor species to present as graded, switch-like, or effectively silent depending strictly on its surface density. When evaluating heterogeneous AC combinatorial drives, the composite expression in A1.20 is bypassed, and the individual AC outputs are summed and passed through the explicit saturable biosensor boundary defined in A1.18.

### Layer 6: Dual-pathway integration architectures

When DRD1 (Gs-coupled) and DRD2 (Gi-coupled) are co-expressed, the total cAMP output is a function of two simultaneous, opposing transducer drives converging at the adenylyl cyclase (AC) layer. To isolate the mechanism of this signal integration, the co-expression models inherit all single-receptor operational building blocks unchanged.

For each pathway *r*∈{*Gs, Gi*}, the receptor occupancy ρ_*r*_, signaling drive *D*_*r*_, and activated transducer fraction *S*_*r*_ remain identical to the single-receptor limits. The maximum possible cAMP output before any Gαi-mediated suppression is applied is defined as the Gαs-driven AC drive:

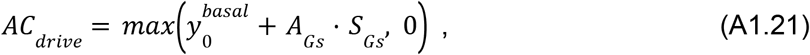

where 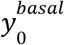 is the intrinsic basal signal, and *A*_*Gs*_ is the maximal cAMP amplitude achievable by the Gαs pathway, bounded by the physical capacity of the cell’s adenylyl cyclase pool.

The maximum capacity of Gαi to suppress this signal is derived from the single-receptor calibration limit. This is formalized as the fractional inhibition limit ε_*max*_ :

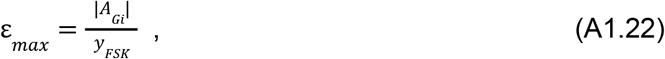

where *A*_*Gi*_ is the maximal negative amplitude driven by DRD2, and *y*_*FSK*_ is the maximally elevated baseline (the forskolin-saturated cAMP ceiling). For the DRD2 pathway, ε_*max*_ approaches 0.68, indicating that maximal Gαi activation can suppress approximately 68% of the total, forskolin-induced cAMP production. To determine how the active Gαi fraction (*S*_*Gi*_) interacts with the Gαs drive, three biological hypotheses are formalized as distinct mathematical models.

#### Model 1: Additive model (homogeneous multiplicative null)

The most parsimonious model assumes that the entire adenylyl cyclase (AC) pool is a single, homogeneous population. Biologically, this implies that both Gαs and Gαi have equal access to all AC molecules, and that Gαi acts as an independent, non-competitive fractional inhibitor of whatever catalytic activity is present. Mathematically, this integration is purely multiplicative:

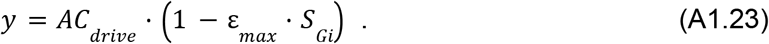

This model establishes the most basic architecture. It assumes no physical interaction between the receptors, no competition within shared G-protein pools, and no isoform-specific compartmentalization at the effector layer. Consequently, it requires zero additional free parameters.

#### Model 2: Cooperative model (receptor-level cooperativity)

The second architecture tests the hypothesis that the Gαs and Gαi pathways interact upstream of the effector layer. Biologically, this represents mechanisms such as active DRD1-DRD2 heterodimerization or synergistic coupling within local G-protein microdomains, mechanisms that the literature frequently, and sometimes prematurely, assumes to be the primary drivers of pathway crosstalk ^14,20^.

Because physical interaction on the two-dimensional plasma membrane is a bimolecular process, the probability of forming a synergistic complex scales with the product of the active receptor densities (the drives, *D*_*Gs*_ and *D*_*Gi*_). This biological assumption is formalized by adding a mass-action synergy term to the null model, with a non-negativity guard to keep the output within physical limits:

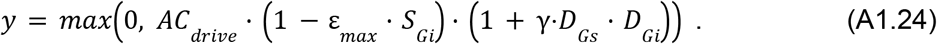

Here, γ is a phenomenological constant that absorbs the theoretical dissociation constant of the heterodimer and its intrinsic signaling efficacy. While this term allows the model to capture non-linear upstream amplification, it lacks microscopic testability.

#### Model 3: AC Integration model (κ-competition)

The third architecture assumes AC isoform heterogeneity in terms of function and expression levels. Mammalian cells express heterogeneous AC pools. As a representative system, CHO-K1 cells specifically express AC6, AC7, and AC9. Biologically, AC6 is canonically whereas AC7 and AC9 are *G*α_*i*_-insensitive. *G*α_*i*_-inhibited,

To respect this physiological constraint, the total cAMP output is partitioned into an isoform-resolved sum, weighted by the transcript fractions (*F*_6_, *F*_7_, *F*_9_). Because AC7 and AC9 are *G*α_*i*_-insensitive (ε_7_ = ε_9_ = 0), the maximal Gαi inhibition must be carried entirely by the AC6 sub-population. To preserve the total cellular calibration, the AC6-specific inhibition term ε_6_ is rescaled:

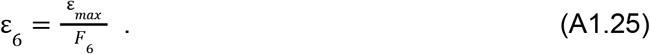

Beyond differential sensitivity, this model assumes stoichiometric competition at the AC6 enzyme. When both pathways are highly active, accumulating *G*α_*s*_ and *G*α_*i*_ compete for allosteric regulation of the AC6 catalytic domains. Consequently, the effective Gαi signal at AC6 is modeled as saturable with respect to the opposing Gαs drive, tuned by a microscopic competition parameter κ:

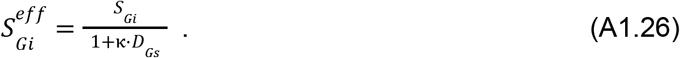

The total cAMP output is therefore the linear sum of the three parallel AC drives, with Gαi inhibition acting exclusively at the competitive AC6 node:

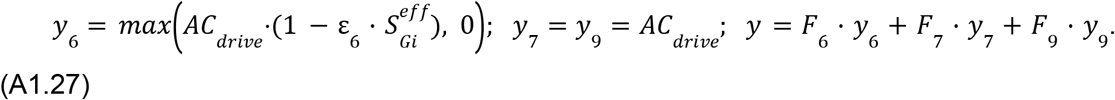

This architecture mathematically enforces the biological assumption that signal integration is determined by the specific effector-isoform composition of the cell, without requiring upstream receptor crosstalk. Crucially, unlike the phenomenological γ parameter, κ is anchored to a specific AC isoform function and is directly testable via targeted depletion of the permissive isoforms.

#### Model 4: Isoform-resolved AC Integration Model (explicit per-isoform signaling functions)

Model 3 couples each receptor’s transducer (Gαs from DRD1, Gαi from DRD2) to a defined pool of three endogenous adenylyl cyclase isoforms (AC6, AC7, AC9). To properly contextualize this model, it is necessary to consider the distinct regulatory families to which these isoforms belong, as they exhibit known biochemical differences under Gαs drive. AC6 (Group III) is Gαs-stimulated and Gαi-inhibited. AC7 (Group II) is Gαs-stimulated, intrinsically Gαi-insensitive, and conditionally potentiated by Gβγ, a mechanism which requires concurrent Gαs activity but can utilize Gβγ released by Gαi-coupled receptors. AC9 (Group IV) exhibits weak Gαs sensitivity and Gαi resistance^22^. A shared mathematical function (as in Figure 5C) would inappropriately force a common Hill coefficient, half-maximal activation, and saturation limit onto enzymes known to differ across all three metrics. For each isoform a separate function can thus be defined:

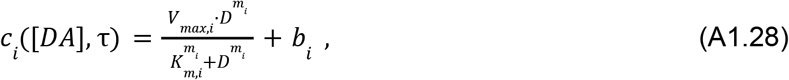

Each parameter carries a physical interpretation: *V*_*max,i*_ is the saturating cAMP amplitude that isoform *i* can produce given an unlimited Gαs supply; *G*,i is the drive at which the isoform reaches half its *V*_*max*_; *m* captures any apparent cooperativity in the isoform’s Gαs activation curve; and *b*_*i*_ is the ligand-independent basal output. The total cellular cAMP response is the *F*_*i*_-weighted sum of these per-isoform contributions, where *F*_*i*_ is the transcript-level fraction of isoform *i* in the AC pool.

